# Host-specific bacterial modulation of airway gene expression and alternative splicing

**DOI:** 10.1101/2025.07.18.665426

**Authors:** Mian Horvath, Hyeon Gu Kang, Te-Chia Wu, Elizabeth Aiken, Diana Cadena Castaneda, Sema Akkurt, Florentina Marches, Olga Anczuków, Karolina Palucka, Julia Oh

## Abstract

The human microbiome varies extensively between individuals. While there are numerous studies investigating the effects of inter-individual differences on microbiome composition, there are few studies investigating inter-individual effects on microbial modulation of the host, or host-specific effects. To address this knowledge gap, we colonized human bronchial epithelial air-liquid interface tissue cultures generated from six different adults with one of three phylogenetically diverse bacteria and compared how each microbe differentially modulated host gene expression in each of the six donors. Microbial treatment had the strongest effect on transcription, followed by donor-specific effects. Gene pathways differed markedly in their donor- and microbe-specificity; interferon expression was highly donor-dependent while transcription of epithelial barrier and antibacterial innate immunity genes were predominantly microbially driven. Moreover, we evaluated whether microbial regulation of alternative splicing was modulated by donor. Strikingly, we found significant non-redundant, donor-specific regulation of alternative splicing exclusively in the Gram-positive commensal microbes. These findings highlight that microbial effects on the human airway epithelium are not only species-specific but also deeply individualized, scoring the importance of host context in shaping microbe-induced transcriptional and splicing responses.

## Introduction

The respiratory epithelium is an essential component of innate immunity, serving as the interface between host immune response and environmental stimuli. A critical factor influencing this interaction is the microbiome, which plays a pivotal role in epithelial physiology(1–3) and host immunity(4–9). Furthermore, the biodiversity present within the respiratory microbiome differentially regulates host-microbiome interactions(9–11). However, while the importance of microbial biodiversity is well recognized, how host factors contribute to differences in these interactions remains poorly characterized and hinders the generalizability of many microbiome findings.

Host variability is a broad concept that encompasses fundamental characteristics such as age and sex, genetic variation, or epigenetic modification from environmental exposures, among others. Regardless of the source of variability, host differences have numerous manifestations. For example, inter-individual variations manifest as differences in immune response(12–14), signaling pathways(15–17), and epithelial barrier(18, 19), all of which modulate the relationship between the epithelium and its microbiome. Furthermore, human genetic studies have highlighted how host genetics affect microbiome composition(19–23) and susceptibility to infectious diseases(17, 18, 24, 25). However, few studies have characterized how inter-individuality affects microbially induced gene expression changes in the epithelium(17, 24).

We previously leveraged 3D air-liquid interface (ALI) tissue cultures, which provide a more physiologically relevant pseudostratified, differentiated cell culture model than traditional monolayers, to examine the transcriptional response of the bronchial epithelium to individual colonization with approximately 60 patient-derived microbes(10). There, we found that microbes differentially activated interferon-stimulated genes, and that interferon stimulation was largely uncoupled from the inflammation response. These responses were microbe-specific and not determined by phylogenetic similarity. Here, we build on this ALI culture system to examine to what degree these responses are donor-versus microbe-specific. We examined transcriptional changes in primary bronchial epithelial cells derived from six adult donors following colonization with one of three representative bacteria, each of which elicited a distinct transcriptional response from the initial screen. We found that the host was a determining factor in interferon stimulation, while changes in antibacterial innate immunity and epithelial barrier genes were primarily microbe-driven. Moreover, we observed a striking microbe-dependent, host-specific differences in alternative splicing, particularly in response to Gram-positive commensal microbes. Taken together, this study provides a detailed evaluation of how host factors modify bacterial modulation of epithelial gene expression and alternative splicing.

## Results

### Generation of 3D primary bronchial epithelial air-liquid interface tissue cultures from different donors

Primary bronchial epithelial cells were obtained from six adult donors (Table S1) and were used to generate air-liquid interface (ALI) tissue cultures as previously described(26–28) (Fig. 1A). ALI culture maturation was monitored using transepithelial electrical resistance (TEER) measurements (Fig. 1B, Table S2). ALI cultures were air-lifted once the TEER exceeded 500 Ω x cm^2^, which indicates the formation of tight junctions. ALI cultures were monitored by bright-field microscopy for beating cilia and mucus formation, markers of epithelial differentiation. Immunofluorescence was used to evaluate cell composition and gross morphology of mature, untreated ALI cultures (Fig. 1C). While there were small differences in relative composition, all donors demonstrated pseudostratified columnar epithelium, and tissues were comprised of all major epithelial cell subtypes: ciliated cells (characterized by acetylated α-tubulin), club cells (SCGB1A1), goblet cells (MUC5AC), and basal cells (CK5).

**Figure 1.**
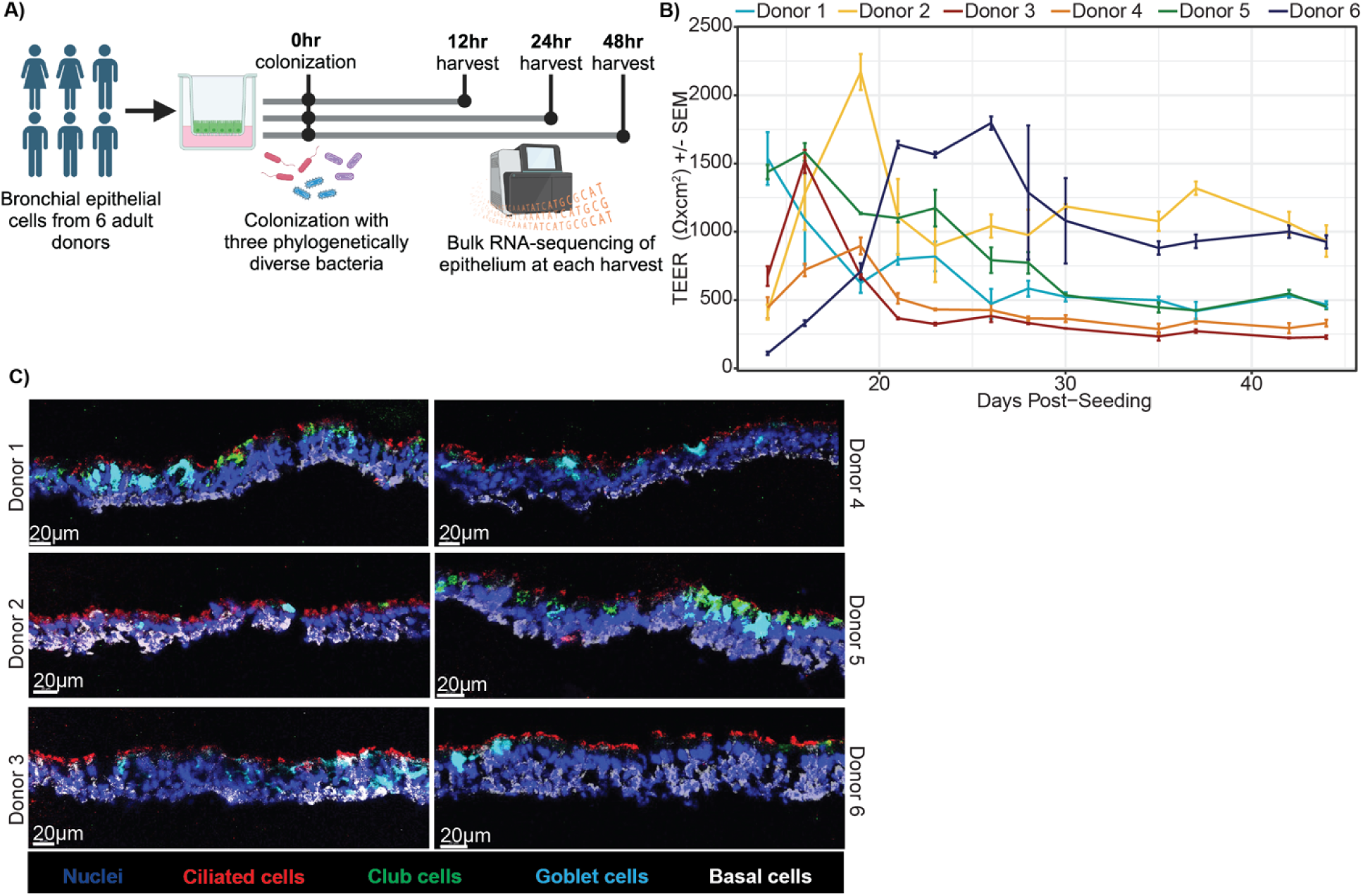
Generation of 3D primary bronchial epithelial air-liquid interface tissue cultures from different donors. **A)** Experimental schematic. **B)** TEER measurements for each donor throughout maturation. Each point is the average of the three ALI measured, and the error bars indicate standard error of the mean (SEM). Lines are colored based on the donor. **C)** Merged cell composition immunofluorescence. Representative immunofluorescence images were obtained from an untreated mature ALI from each donor stained for nuclei (DAPI, blue), ciliated cells (acetylated α-tubulin, red), Club cells (SCGB1A1, green), goblet cells (MUC5AC, cyan), and basal cells (CK5, white). Scale bar, in white in lower left corner, is 20μm. n = 1.

### Microbial effects on host transcription differ by donor and vary temporally

Mature ALI cultures were colonized with 10^7^ CFUs (colony forming units, a measure of live bacteria) of one of three previously characterized strains of bacteria: *Klebsiella* (*K.*) *aerogenes*(10), *Rothia* (*R.*) *aeria*(10), or GFP-tagged *Staphylococcus* (*S.*) *epidermidis* (Tü3298)(10, 29), or vehicle for 12, 24, and 48 hours (Fig. 1A). These microbes were selected as genetically diverse representatives of differential interferon and inflammatory response in our previous screen. Specifically, *K. aerogenes* is a Gram-negative, motile opportunistic pathogen; *R. aeria* is a Gram-positive commensal; and *S. epidermidis* is a Gram-positive commensal with opportunistic potential. At each time point following colonization, the apical surface wash was plated for CFUs to approximate microbial growth (Fig. S1, Table S3) and total RNA from the host epithelial cells was extracted and sequenced. Microbial growth was highly variable between microbes, but no gross host effects were observed (Kruskal-Wallis test, P-value > 0.05) except for *K. aerogenes* at 12 hours. We also excluded 24 genes whose transcription were determined to be driven by CFUs alone based on a linear mixed effects model implemented in MaAsLin2(30), as we previously performed(10).

We first examined overarching transcriptional similarities between donors. Principal components analysis (PCA) for each time point showed that while neither microbial treatment nor donor were strong contributors to sample variability at 12 hours, by 24 hours (and subsequently at 48 hours) microbial treatment was the strongest contributor (Fig. 2A). *S. epidermidis* and *R. aeria* elicited a more similar effect on the host vs. *K. aerogenes*, potentially due to their more commensal nature. Host-specific differences (as measured by centroid distance) also accumulated over time, with minimal separation between donors and low variance explained at 12 hours, peaking at 24 hours, then declining by 48 hours. This suggested that there is a universal early and late response to microbial colonization, with the most host-specific variability occurring at an intermediate time point. Interestingly, the separation seen at 24 hours corresponded with donor sex and age, suggesting that sex and age are strong factors driving transcriptome-wide, inter-individual variability.

**Figure 2.**
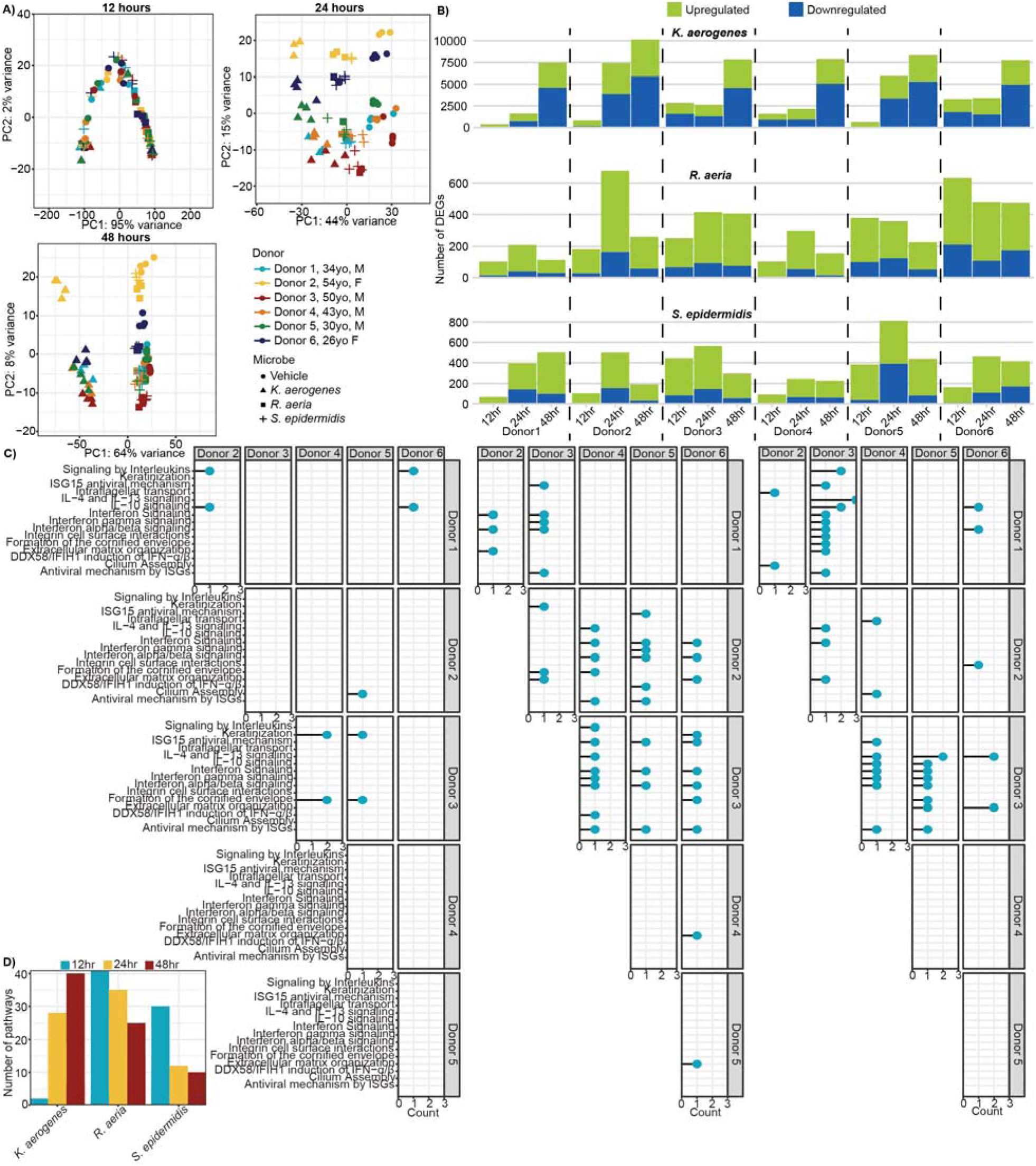
Microbial effects on host transcription differ by donor and vary temporally. **A)** Whole transcriptome principal component analysis (PCA) for each time point. Points are shaped based on the treatment and colored based on the donor. Donors are labeled with their age (yo = years old) and sex. n=3-4 ALI cultures. **B)** Number of differentially expressed genes (DEGs) for each microbial treatment relative to vehicle control for each donor at each time point. DEGs were defined as adjusted P-value < 0.05 and absolute log_2_ fold change > 1. The bottom blue bar represents downregulated genes, and the upper green bar represents upregulated genes. Dotted lines separate each donor. **C)** Reactome enrichment analysis of the genes differentially modulated between pairwise donor comparisons in response to microbial colonization for each time point. The top 15 pathways with the lowest adjusted P-values across all time points are visualized. X-axis and Y-axis labels depict the donors compared in each pairwise comparison. X-axis quantifies the number of microbial treatments for which this gene set was significantly enriched. Empty plots were ones in which there were no significantly enriched gene sets. IFN = interferon; IL = Interleukin; ISG = Interferon-stimulated gene. **D)** Total number of significantly enriched pathways for each microbial treatment (x-axis) at each time point (bar color).

To further compare gross differences in host response to microbial colonization, we quantified the number of differentially expressed genes (DEGs) for each microbial treatment at each time point, per donor (Fig. 2B, Table S4). Unsurprisingly, *K. aerogenes* was the most modulatory regardless of donor with ∼10x as many DEGs at most time points, suggesting this pathogen causes a strong disruption to host homeostasis. However, host responsiveness varied extensively across microbial treatments and also differed temporally. For example, donor 2 had the most DEGs at 24 hours when treated with *K. aerogenes* or *R. aeria,* but not *S. epidermidis* for whom donor 5 experienced the strongest response at 24 hours. These results highlighted a surprising specificity to the host-microbe interactions.

Finally, to generalize if there were common functional pathways that differed between donors in response to microbial colonization, we pairwise compared enriched Reactome pathways from genes differentially expressed between donors in response to microbial treatment (Fig. 2C, Table S5). We found that the early response was relatively common between donors/microbes, with host- and microbe-specific responses appearing at later timepoints. For example, while there were few early (12 hour) enriched pathways, by 24 hours proinflammatory pathways, antiviral interferon pathways, and epithelial barrier pathways were enriched, but typically only by a single microbe. By 48 hours, many more gene sets were enriched across microbes. Despite *K. aerogenes* stimulating a stronger host response than the other two microbes, the enriched pathways were not exclusive to *K. aerogenes* (Fig. 2D). In fact, at 12 and 24 hours, treatment with *R. aeria* resulted in more enriched pathways than either of the other microbes, suggesting the strength of the host response does not predict the degree of similarity between hosts. Altogether, microbe-specific factors play a role in determining host specificity, and temporality may contribute to the degree of microbe and donor specificity.

### Stimulation of interferon is microbe- and donor-specific, but antibacterial innate immunity and epithelial barrier is predominantly microbe-driven

The enriched pathways consisted of immune pathways (interleukins and interferons) and epithelial pathways (keratinization, cilium, and cornified envelope), suggesting that these were donor-modulated functions. Thus, we further examined variations in epithelial barrier genes and innate immune activation. We manually curated epithelial barrier gene lists (Table S6) to include: mucins which comprise mucus(31) (Fig. S2), keratins which provide structural integrity(31) (Fig. S3), cilia(32) (Fig. S4), and apical junctions (junctions)(33) (Fig. S5). To quantify and generalize the activation of epithelial barrier genes, we generated a ‘score’ for these categories represented by the median log_2_ fold change of the genes in each gene list(10) (Fig. 3A, Table S7). Generally, transcription of epithelial barrier genes was primarily microbe-dependent, with *K. aerogenes* inducing the largest effect. However, there was limited donor-specificity. For example, at 24 and 48 hours, donor 2 had an outsized response compared to other donors irrespective of microbe. Donors 1 and 3 appeared to have a delayed response to *K. aerogenes*, with mild changes in gene scores (excluding ciliary score) until 48 hours, while all other donors had a stronger response by 24 hours.

**Figure 3.**
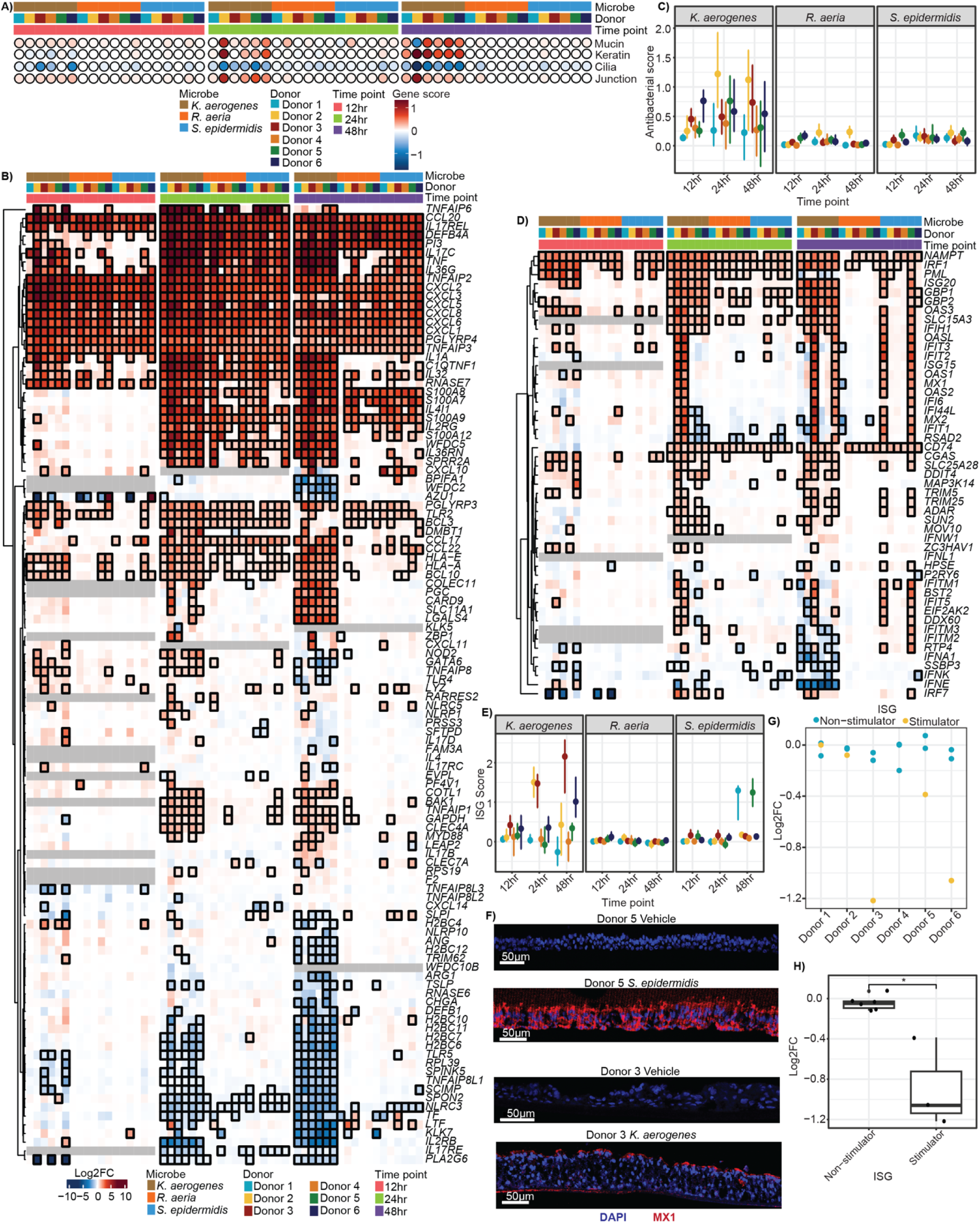
Stimulation of interferon is microbe- and donor-specific, but antibacterial innate immunity and epithelial barrier is predominantly microbe-driven. **A)** Dotplot of epithelial barrier gene lists where each row is a different gene list and each column a different sample. Samples are annotated by color for their microbial treatment, donor, and time point. Points are colored based on their gene score (median log_2_ fold change (FC) over vehicle control). **B)** Heatmap of transcriptional response in a manually curated list of antibacterial innate immunity (non-interferon cytokines and antimicrobial peptides). Each row represents a different gene and each column a different sample. Columns were hierarchically clustered. Each column was annotated by color with the sample’s microbial treatment, donor cells, and time point. Cells were colored by the log_2_FC relative to the appropriate vehicle control. Outlined boxes have an adjusted P-value < 0.05. Gray boxes represent genes that were filtered out due to low gene counts at a specific time point. Genes that were not present in at least 2/3 time points were removed. **C)** Dotplot of the antibacterial score for each sample. Points are colored by donor according to the legend in Fig. 3B. Error bars represent the 95^th^ confidence interval determined by bootstrapping. **D)** Heatmap of interferon stimulated genes (ISGs) relative to vehicle control as in Fig. 3B. **E)** Dotplot of the ISG score for each sample as in Fig. 3D. **F)** Representative immunofluorescence images of ISG MX1 (red) and DAPI/nuclei (blue) following 48 hours of colonization. n = 3 ALI cultures. **G)** Dotplot of the log_2_FC for *PIAS3* expression at 12 hours. Each point represents a different microbial treatment, and points were colored yellow if that microbe-donor pairing resulted in ISG expression at 24 or 48 hours. **H)** Boxplot of the log_2_ fold changes for donors 3, 5, and 6. Wilcoxon test was used to compare between ISG stimulators and non-stimulators. For relevant plots, * represented P-value < 0.05, ** P-value < 0.01, *** P-value < 0.001, and ns = non-significant. For boxplots, box middle represents the median, box edges represent 25^th^ and 75^th^ quartiles, and outlier values are separate points.

Antibacterial genes, *i.e.*, non-interferon cytokines and antimicrobial peptides(34, 35) also had less pronounced donor variability (Fig. 3B, Table S6). For example, genes associated with antibacterial innate immune response such as immune cell chemotaxis(36–38) and antimicrobial peptides(39) were upregulated regardless of timepoint, donor, or microbial treatment, and changes in other genes were primarily driven by *K. aerogenes*. Some donor-specific effects were observed; for example, donor 1 had a relatively weak response across microbial treatments/time and conversely, once again donor 2 had an outsized response and donor 6 also seemed to have a stronger response, albeit much weaker than donor 2. While some of the other microbial treatments appeared to have enhanced activation of antibacterial innate immunity, their gene scores were nominal, likely because of the broadness of the antibacterial innate immunity gene set (Fig. 3C, Table S7).

We then examined interferon stimulated genes (ISGs)(40), whose induction was notably microbe-specific in our initial screen. Stimulation of ISGs, both at the gene level (Fig. 3D, Table S6) and as an aggregated score (Fig. 3E, Table S7), was highly microbe- and donor-specific. *K. aerogenes* and *S. epidermidis* stimulated ISG expression in a subset of donors. Strikingly, they did not stimulate gene expression in the same donors, indicating that ISG induction is not entirely driven by the donor but rather a synergy between donor and microbe. Finally, to confirm that bacterial treatment was stimulating expression of antiviral innate immune genes, we conducted immunofluorescence of MX1, an ISG with direct antiviral function (Fig. 3F).

We wanted to confirm that the observed variability in host responses was due to donor-specificity and not confounding factors such as hypoxia(33, 41) (Fig. S6, Table S6) or apoptosis(42, 43) (Fig. S7, Table S6). Unsurprisingly, upregulation of both pathways was exclusive to *K. aerogenes* colonization, which was also the most vigorous proliferator. This may also explain the enhanced antibacterial innate immune response and changes in epithelial barrier genes that were seen with *K. aerogenes* colonization. However, given that there were no statistically significant differences in growth between donors for *K. aerogenes* or the other microbes, differential growth would not fully explain the donor-specific differences observed, in particular the differential stimulation of interferon. Furthermore, ISG stimulators did not have a statistically significant increase in the hypoxic markers *HIF1A* (Wilcoxon test, P-value = 0.65) and *EPAS1* (*HIF2A*), which was actually slightly repressed with ISG stimulators (Wilcoxon test, P-value 0.0024).

Given that interferon is constitutively expressed at low levels in the epithelium, we were curious if basal interferon expression contributed to the observed donor- and microbe-specificity (Fig. S8, Fig S8). While baseline ISG expression (vehicle) varied between donors, it did not correlate with whether ISG expression was observed following microbial colonization. Furthermore, the basal expression level was far lower than that seen with microbial stimulation, disproving that differential expression of ISGs was due to basal interferon already being at maximum capacity.

Finally, although we did not anticipate that the mechanisms of bacterial stimulation of interferon would be conserved between the Gram-positive *S. epidermidis* and Gram-negative *K. aerogenes*, nor the causes of differential host responses to be shared between our donors, we sought to identify potential sources of differential interferon stimulation. First, we queried interferon pathway genes such as pattern recognition receptors, required transcription factors, and negative regulations; however, we found no correlations between their expression and interferon stimulation. To broaden our search, we identified any DEGs that were shared between either ISG non-stimulation conditions (donor-microbe pairing) or ISG stimulation conditions and then filtered for genes with known connections to interferon. We identified one gene, *PIAS3*, that correlated with ISG stimulation across multiple donors and both microbes at 12 hours, a time point preceding ISG stimulation (Fig. 3G-H). *PIAS3* has been shown to covalently attach SUMO-1 (a ubiquitin-like post-translational modification) to IRF1 and STAT3, repressing the activity of both proteins(44, 45), which are both implicated in the expression of interferon and ISGs(46–49). Additionally, *PIAS3* can inhibit STAT3’s DNA-binding site, further inhibiting its function(50). At 12 hours, we observed a statistically significant decrease in *PIAS3* expression only in ISG stimulating donor-microbe pairs for donors 3, 5, and 6 (Fig. 3H-G, Wilcoxon test, P-value = 0.024). We hypothesize that these donors are susceptible to bacterial modulation of *PIAS3* expression due to either genetic or epigenetic variation and that the subsequent host-microbe interactions decreased the expression of *PIAS3,* relieving the normal inhibition present in the absence of viral infection, thus contributing to the expression of interferon and ISGs.

### Alternative splicing in response to microbial colonization is host- and microbe-dependent

Thus far, we demonstrated that microbial colonization modulates host gene expression in a host-dependent manner; however, gene expression alone does not fully predict protein expression. An additional layer of regulation is provided by alternative splicing, in which a single gene can produce multiple transcripts and protein isoforms. Indeed, the lack of genes whose expression correlated with the highly host-specific ISG stimulation suggested to us that additional mechanisms beyond gene expression may account for some of these observed host-specific differences. While a number of extrinsic factors regulate alternative splicing, only a few studies have investigated microbial regulation of alternative splicing(51–54), and even fewer have done so within the context of extracellular bacterial colonization(55). Thus, to determine if microbial colonization affects alternative splicing in a host-dependent manner, we predicted splicing isoforms following 48 hours of colonization. We excluded two donors from the analysis with *K. aerogenes* due to insufficient read counts.

First, we examined if microbial colonization induced alternative splicing relative to vehicle treatment, identifying differential splicing events for each donor-microbe pairing. Cassette exons (skipped exons) were the most common splicing event across all donors and microbes, followed by mutually exclusive exons (Fig. S9, Table S8). For gross comparison, we defined differentially spliced genes (DSGs) across all donors and microbial treatments as any gene with at least one differential splicing event (Fig. 4A-B, Table S9). Paralleling transcriptional data, *K. aerogenes* induced more DSGs than the commensal microbes. When we compared the number of genes that were differentially expressed and/or differentially spliced, we found the total genes modulated by *K. aerogenes* were approximately split as DEGs or DSGs. Strikingly, despite having a modest effect on host gene expression, *S. epidermidis* and *R. aeria* had a profound effect on splicing with approximately one third as many DSGs as *K. aerogenes* (in contrast to the ∼1/10 DEGs). This is even more profound when contrasted with their bioburden, with *K. aerogenes* achieving significantly higher CFUs than the commensal microbes. Furthermore, the commensal microbes’ DSGs made up more than 90% of the total number of genes modulated by *S. epidermidis* or *R. aeria* colonization. This suggests that while these microbes may not be inducing overt gene expression changes, they are exerting an active function on the respiratory epithelium.

**Figure 4.**
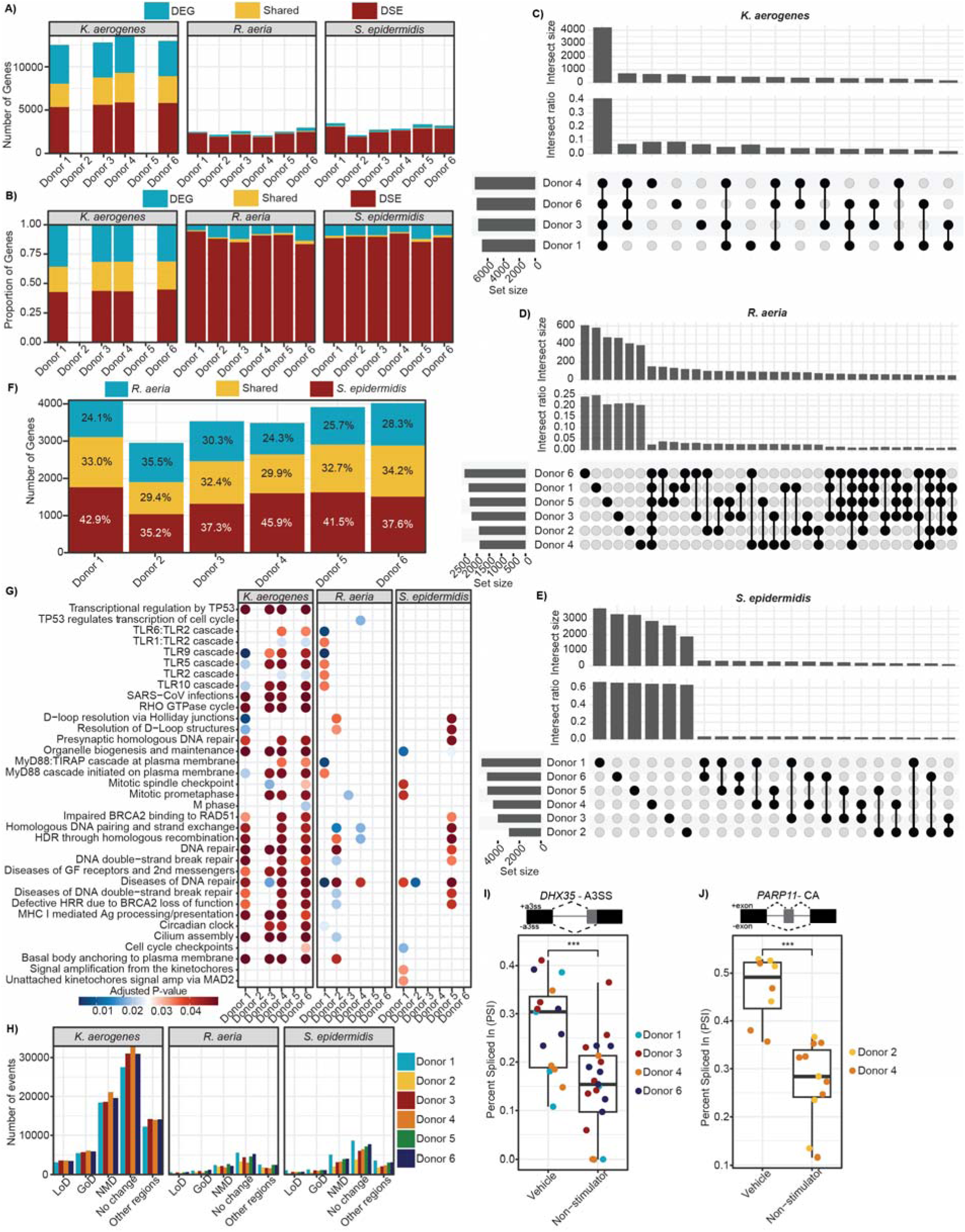
Alternative splicing in response to microbial colonization is host- and microbe-dependent. **A)** Number of differentially spliced genes (DSGs, red), DEGs (blue), or genes that were both differentially spliced and differentially expressed (shared, yellow) for each microbe-donor pairing relative to vehicle control. Donors 2 and 5 colonized with *K. aerogenes* were excluded from analysis due to insufficient read counts. **B)** Proportion of DSGs, DEGs, and shared genes from Fig. 4A. **C)** Upset plot of DSGs for each donor when colonized with *K. aerogenes*. Set size represents the total number of DSGs. Donors are sorted by decreasing set size. Intersect size represents the number of DSGs shared between the indicated donors (black points connected by lines). Only intersections with a size of at least 50 are displayed. Intersect ratio normalizes the intersect size by union size (all unique elements across the sets). **D)** Upset plot for DSGs following colonization with *R. aeria*. **E)** Upset plot for DSGs following colonization with *S. epidermidis*. **F)** Number of DSGs that were unique to colonization with *R. aeria* (blue), *S. epidermidis* (red), or shared between the two (yellow). The percent of each component is labeled on the bar. **G)** Reactome pathway enrichment analysis of the DSGs for each microbe-donor pairing. The top 25 pathways with the lowest adjusted P-value are visualized. The points are colored by adjusted P-value. Non-significant pathways (P-value > 0.05) were not visualized. GF = growth factors; 2^nd^ messengers = secondary messengers; Ag = antigen. **H)** Number of splicing events predicted to result in gain of domain (GoD), loss of domain (LoD), nonsense mediated decay targets (NMD), no change, and changes in other transcript regions shown for each microbe and colored by donor. **I)** Inclusion of an alternative 3’ splice site in *DHX35* is quantified sing RNA-seq data as percent spliced in (PSI) value for indicated samples for which there was predicted to be a domain change. Above the boxplot is a diagram of the splicing event, where the black box represents the upstream and downstream exons, and the gray box represents the alternative splice sites. +a3ss represents when the alternative 3’ splice site is included and -a3ss represents when the alternative 3’ splice site is excluded. Samples are organized by treatment with vehicle (Vehicle) and non-ISG stimulator microbe (Non-stimulator). Significance was calculated using Wilcoxon test. **J)** Inclusion of a cassette exon (skipped exon) in *PARP11* is quantified using RNA-seq data as percent spliced in (PSI) for the indicated samples for which there was predicted to be a domain change. Above the boxplot is a diagram of the splicing event, where the black box represents the upstream and downstream exons, and the gray box represents the cassette exon. +exon represents when the cassette exon is included and -exon represents when the cassette exon is excluded. Samples are organized by treatment with vehicle (Vehicle) and non-ISG stimulator microbe (Non-stimulator). Significance was calculated using Wilcoxon test. For relevant plots, * represented P-value < 0.05, ** P-value < 0.01, *** P-value < 0.001, and ns = non-significant. For boxplots, box middle represents the median, box edges represent 25^th^ and 75^th^ quartiles, and outlier values are separate points.

While the number of DSGs were generally comparable between donors for each microbial treatment, we were curious if each microbe was inducing splicing events in the same genes. Strikingly, we found a large difference between pathogen and commensal species. Approximately 40% of the DSGs associated with *K. aerogenes* colonization were shared between all four donors, and less than 10% of DSGs were unique to a donor (Fig. 4C). This is contrast to *R. aeria* (Fig. 4D) and *S. epidermidis* (Fig. 4E), for which only ∼2.5% of DSGs were shared between all six donors (∼25% were unique to a donor) and only an additional 5% of DSGs shared between the same four donors as *K. aerogenes*. Notably, there was only partial overlap in DSGs for *S. epidermidis* and *R. aeria* – for each donor, almost a third of the DSGs were shared between the two microbes, with a quarter unique to *R. aeria* and almost half unique to *S. epidermidis* (Fig. 4F). This pattern is further recapitulated when we compared similarity between differential splicing events (Fig. S10). Unsurprisingly the proportion of shared splicing events across donors was significantly lower; ∼60% of *R. aeria* and *S. epidermidis*’ splicing events were unique to each donor, compared to ∼40% for *K. aerogenes*, despite analyzing fewer donors for the latter. This suggests that not only does *R. aeria* and *S. epidermidis* have a more pronounced effect on alternative splicing than they do on gene expression, their effect is highly host-specific and non-redundant (*i.e.*, they induce splicing isoforms in different genes).

We then examined potential functional consequences of these splicing events. Reactome pathway enrichment for the DSGs showed more similar pathway enrichment for *K. aerogenes,* but more distinct differences for *R. aeria* and *S. epidermidis* (Fig. 4G, Table S10). Some pathways were almost universally enriched, such as diseases of DNA repair. However, most pathways were unique to a few microbe-donor pairings, with few universally conserved across donors or microbes. Corresponding to our earlier prediction that alternative splicing may contribute to differential ISG stimulation, many toll-like receptor pathways (TLR and Myd88; responsible for pathogen recognition and initiation of immunity) were enriched.

We then used SpliceDecoder(56), a computational pipeline to predict the functional consequences of each splicing event, such as whether it would result in a change to an annotated protein domain leading to gain or loss of a domain, introduce a premature termination codon that would target the transcript for nonsense-mediated mRNA decay (NDM), or alter other regions including 5’ or 3’ untranslated regions (Fig. 4H, Table S11). Notably, *K. aerogenes* was associated with a high rate of transcripts predicted to undergo NMD across all four donors, which might explain why it has a higher rate of gene downregulated in comparison to the other microbes (Fig. 2A), since NMD is a mechanism of physiological gene downregulation(57). Despite differences between microbes, the rates of each functional consequence were relatively consistent across donors for each microbe.

Out of the genes predicted to have a splicing event leading to a gain or loss of protein domain, we identified two genes that correlated with differential ISG expression, *DHX35* and *PARP11*. *DHX35* is a co-sensor for *RIG-I*, a pattern recognition receptor that stimulates expression of interferon(58). For some of the ISG non-stimulators in donors 1, 3, 4, and 6, we predicted there would be a statistically significant decrease in a *DHX35* splicing event using an alternative 3’ splice site, which was predicted to disrupt the location of the start codon, adding 486 nucleotides (162 amino acids) to the N-terminus (Helicase ATP-binding domain). Therefore, we predicted increased expression of a protein isoform with a truncated N-terminus relative to the vehicle treated samples (Wilcoxon test, P-value = 1.6 x 10^-3^, Fig. 4I, Fig. S11A).

*PARP11* regulates expression of *IFNAR1*, the receptor for Type 1 interferon, targeting it for degradation(59). For some of the ISG non-stimulators in donors 2 and 4, we predicted there would be a statistically significant decrease in the inclusion of an alternative exon, which would result in the addition of 102 nucleotides (34 amino acids) to the WWE domain. This would result in a protein isoform with a modified WWE domain relative to the vehicle treated samples (Wilcoxon test, P-value = 5.3 x 10^-5^, Fig. 4I, Fig. S11B). While the effects of neither splicing event (*DHX35* and *PARP11*) are characterized, these changes are predicted to impact the protein folding by AlphaFold and may modulate their activity (Fig. S12). Taken together, our results show extensive variability in host- and microbe-specific effects on alternative splicing in addition to transcriptional differences, showing a surprising variability in the host-microbiome interactions.

## Discussion

In summary, we present evidence for host-specific determinants of transcriptional response to microbial colonization. These could include numerous potential sources, including genetic variability such as allelic variation, single nucleotide variants, and larger structural variants. Adding another layer of complexity, recent studies have demonstrated that epithelial cells can develop a “memory” to environmental exposures through epigenetic modifications(60–62). The host-specificity observed in our study may result from any of the aforementioned factors either individually or in concert.

All of the microbes selected for this study induced host-specific responses in innate immunity and epithelial barrier genes, with the most pronounced host-specificity observed in interferon stimulation. Interferon is the canonical antiviral innate immune response, although studies have shown interactions between extracellular bacteria and interferon(10, 63–74). We previously observed extensive differences in the ability of phylogenetically diverse microbes to stimulate interferon in a single donor(10), which was also observed here in addition to host-specificity. *K. aerogenes* only stimulated interferon in donors 2, 3, and 6, and *S. epidermidis* only stimulated interferon in donors 1 and 5. Furthermore, donor 4 did not express ISGs in response to either microbe, despite expressing ISGs in response to viral infection(28). This demonstrates that while both microbes are capable of stimulating interferon, it is not universal across donors and is thus likely multifactorial and driven by both to-be-determined host and microbial factors.

The multifactorial nature of microbial stimulation of ISGs complicates prediction of genes underlying differential interferon stimulation. We conjecture that *S. epidermidis* and *K. aerogenes* may have distinct mechanisms for stimulating interferon based on their genetic variability, which is further supported by the several known bacterial mechanisms to stimulate host interferon demonstrated in peripheral blood mononuclear cells and epithelium(66, 71, 75–79). Similarly, host factors may modulate microbial stimulation of interferon that could also be distinct between donors based on the complex patterns of activation. Another challenge in identifying the causes of host variability is our reliance on transcriptional data, which does not capture post-translational modifications – such as phosphorylation, which is critical for signaling pathways like interferon; acetylation, which can be regulated by microbes(80, 81); and sumoylation, regulated by *PIAS3*, a gene we found to be differentially expressed between ISG stimulators and non-stimulators – as well as alternative splicing.

Alternative splicing was strikingly microbe- and donor-specific and did not mirror patterns seen in the transcriptional data. In particular, while colonization with the pathogenic *K. aerogenes* led to significantly more DEGs than *R. aeria or S. epidermidis*; the commensal microbes induced a more comparable number of DSGs. While DSGs accounted for approximately half of the genes modulated by *K. aerogenes*, more than 90% of the genes modulated by *R. aeria* and *S. epidermidis* were alternatively spliced. Even accounting for the smaller number of donors analyzed for *K. aerogenes*, *R.aeria* and *S. epidermidis* still showed far more donor-specific splicing: 60% of *K. aerogenes*’ DSGs were shared across all four donors, while fewer than 10% of the commensals’ DSGs overlapped with those same donors – a pattern that was recapitulated when examining shared splicing events. This may reflect a key difference in microbial nature, with pathogenic microbes stimulating broad and aggressive gene expression changes in the host for infection control, while commensal microbes drive more subtle but still significant regulation through alternative splicing. Furthermore, we observed that only one third of the DSGs were shared between *S. epidermidis* and *R. aeria*, suggesting that not only are they predominantly exerting their effect through alternative splicing but that their effects are non-redundant, although it is unknown if together the effects would be synergistic or antagonistic.

When we predicted the functional consequence of the splicing events, we noticed cases where splicing patterns could help to explain the observed transcriptional changes. For example, colonization with *K. aerogenes* resulted in a high number of spliced transcripts predicted to be nonsense-mediated decay targets, which corresponds with *K. aerogenes*’ high number of downregulated genes, since nonsense-mediated decay can contribute to regulation of gene expression(57). We also predicted functional domain changes in *DDX53* and *PARP11*, genes that are known to interact with interferon signaling and may contribute to differential stimulation of interferon. The changes in splicing patterns observed provide a compelling example of how microbiome-driven gene expression alterations extend beyond transcription to influence the functional repertoire of proteins.

Taken together, our results underscore the complex, multifaceted nature of microbiome regulation of host gene expression, suggesting that modifications in both transcription and alternative splicing contribute to a host’s response to microbiome colonization. Our findings exhort that studies of host-microbiome interactions must take into account host genetic variability before generalizing microbial functions on their host.

## Methods

### Air-liquid interface cell culture (ALI) cultivation

Deidentified human lung tissue was obtained from NDRI (Project: RPAK1 01) under the approval of the Human Subjects Institutional Review Board at The Jackson Laboratory and were used in the context of the U19AI142733 grant at the Jackson Laboratory.

Air-liquid interface (ALI) cultures were generated as previously described(26–28). Briefly, cryopreserved lung tissue (FBS with 10% DMSO) were thawed in Advanced DMEM/F12 media containing 1x Penicillin/Streptomycin, 10mM HEPES, and 1x GlutaMAX (AdDF+) and digested in 10mL of AdDF+ containing 1-2mg/mL of collagenase I for 1 hour shaking on an orbital shaker at 37°C and 5% CO_2_. Following digestion, the tissue was sheared with sterile slides and strained through a 100μm filter into a single cell solution. The cell solution was then counted, centrifuged (300xg for 5 min), and resuspended in AdDF+ supplemented with 500ng/mL R-Spondin 1, 25ng/mL FGF-7, 100ng/mL FGF-10, 100ng/mL Noggin, 1X B-27 supplement, 50μg/mL Primocin, 500nM A83-01, 5μM ROCK inhibitor Y-27632, 5μM/mL SB202190, 1.25mM/mL N-Aceylcysteine, and 5mM Nicotinamide (AO media) with 10mg/mL Matrigel (Cultrex growth factor reduced BME type 2; Trevigen #3533-010-02). The cell suspension was dispensed at 300,000 cells/40uL per well of a pre-warmed 24-well plate. Once gellification occurred, the organoid cultures were kept in 400uL of AO media with daily media changes. Organoid cultures were passaged 4-7 times and then dissociated into a single-cell stock of primary lung organoid-derived epithelial cells. The cell stocks were then cryopreserved.

To generate air-liquid interface (ALI) cultures, lung organoid-derived epithelial cells were seeded onto 24-well transwell inserts with 0.4μm pore size (Corning #3470) coated with 30μg/mL collagen I (Collagen I Rat Protein, Tail; Gibco #A1048301) at 30,000 cells/100μL per well. Once the cells were seeded, the ALI cultures were grown according to the manufacturer’s instructions (STEMCELL Technologies):

1. Maintenance in Pneumacult-Ex Plus Medium (STEMCELL Technologies #05040) containing 10μM ROCK inhibitor Y-27632, 96ng/mL hydrocortisone, and 1x Penicillin-Streptomycin until 100% confluent (8-12 days). Media was changed every 2-3 days with 100μL in apical chamber and 500μL basal chamber.
2. ALI culture differentiation in Pneumacult-ALI Medium (STEMCELL Technologies #05001) containing 4μg/mL heparin, 0.48μg/mL hydrocortisone, and 1x Penicillin-Streptomycin (ALI+ media) and 10μM ROCK inhibitor Y-27632 (5-7 days). Media was changed every 2-3 days with 100μL in apical chamber and 500μL basal chamber. Transepithelial electrical resistance (TEER) was measured with every media change until TEER values > 500Ω/cm^2^.
3. ALI culture differentiation in ALI+ media without Rock inhibitor Y-27632 (28 days). ALI cultures were air-lifted (media removed from apical chamber) and media in the basal chamber (500μL) was changed every 2-3 days for an additional 28 days. ALI cultures were checked under brightfield microscope for cilia beating (approximately by 2 weeks) and mucus formation (by 2-4 weeks). TEER measurements continued until maturation.

### Immunofluorescence staining

ALI cultures were embedded in optimal cutting tissue compound (OCT) and snap-frozen at -80°C. Frozen sections were cut at 8 µm, air dried on Superfrost plus slides, fixed with 4% paraformaldehyde for 15 minutes, then permeabilized with 1X PBS/0.1% Triton-X for 15 min. Tissue sections were treated with Fc Receptor Block (#NB309, Innovex Bioscience) for 40 minutes followed by Background Buster (#NB306, Innovex Bioscience) for 30 minutes. The sections were stained with either anti-MX1 primary antibody (polyclonal Rabbit N2C2, #GTX110256, GeneTex) or anti-SCGBA1 (Clone 394324, R&D) and anti-acetylated-alpha-tubulin (Clone 6-11B-1, Thermofisher) for one hour followed by the appropriate secondary antibody for 30 min in 1X PBS/5% BSA/0.05% saponin. For cell composition staining, tissues were washed, and secondary antibodies were saturated using mouse normal serum diluted at 1/20 in 1X PBS for 15 minutes. Then, sections were stained with directly conjugated anti-MUC5AC AF700 (Clone45M1, Novus) and anti-cytokeratin 5 AF594 (Rabbit Polyclonal, Novus) for 1 hour and washed. Finally, sections were counterstained with 4’,6-diamidino-2-phenylindole (DAPI) then, mounted with Fluoromount-G (#00-4958-02, Thermo Fisher Scientific), acquired using a Leica SP8 confocal microscope (Leica Microsystems) for high resolution images and analyzed using Imaris software (Bitplane, Oxford Instruments). Following cutting, all staining techniques were performed at room temperature.

### Transepithelial electrical resistance (TEER) measurements

TEER measurements were taken using EVOM2 Epithelial Volt/Ohm TEER meter (World Precision Instruments) and STX2 Electrode (World Precision Instruments) every 2-5 days starting when ALI cultures were switched to Pneumacult-ALI Medium and ending when ALI cultures had finished differentiating. To take readings, ∼200uL of media was added to the ALI culture’s apical compartment. Prior to readings and in between each ALI culture, electrodes were sterilized twice in 70% ethanol and rinsed twice in dPBS. At each TEER measurement, three ALI cultures from each donor were measured. The same three ALI cultures were measured at each time point. In between uses, electrodes were stored in 0.1 M NaCl.

### ALI Treatment

The bacteria used for colonization(10, 29) were grown from single colonies overnight in sterile 1x TSB with 0.1mg Vitamin K and 5mg heme / 1L. Bacteria were washed once with sterile antibiotic-free Pneumacult-ALI Medium before ODs were obtained by measuring absorbance (600nM) in semi-microcuvettes using Eppendorf BioSpectrometer. Bacteria were resuspended in sterile antibiotic-free Pneumacult-ALI Medium with a final concentration of 10^7^ CFUs per 25µL.

ALI cultures were treated on Day 30-33 post-airlift (some donors required airlifting earlier than other donors). 1.5 weeks prior to treatment, ALI cultures were switched to antibiotic free media. Due to the lack of physiological mucus clearance in this *in vitro* model, just prior to treatment excess mucus was washed off of the ALI cultures by adding 100µL of sterile dPBS to the apical chamber. ALI cultures were incubated with dBPS for 10-15 minutes at 37°C. This was repeated for a total of 2 washes. Following the washes, the ALI cultures were then dosed with 25uL of microbial isolate (10^7^ CFUs) or vehicle (sterile antibiotic-free Pneumacult-ALI Medium). Following treatment, ALI cultures were incubated for 12, 24, or 48 hours prior to harvest. Extra inoculum was serially diluted in sterile dPBS and grown on tryptic soy agar (TSA) plates to quantify the number of microbes added to the ALI cultures.

At harvest, 200µL of transepithelial electrical resistance buffer was added to the apical chamber of each ALI culture, pipette mixed 3x, and removed. ALI cultures were cut in half with a scalpel. One half was dissolved in 140µL of Buffer RLT (Qiagen) + 1% beta-mercaptoethanol and stored at -80°C. The other half was embedded in OCT, snap-frozen, and stored at -80°C. The apical wash was serially diluted in dPBS and plated on TSA plates to quantify CFUs at time of harvest. Basal media was plated on TSA plates to determine bacterial contamination of basal compartment. Any ALI cultures with bacterial contamination of the basal media were excluded from analysis, resulting in 3-4 biological replicates per treatment.

### RNA extraction and RNA-seq

RNA extraction and sequencing library preparation were performed in a sterile tissue culture hood. RNA was extracted according to manufacturer’s instructions using the RNeasy 96 QIAcube HT kit, eluted into nuclease-free water, and stored at -80°C until sequencing preparation. RNA quality was evaluated using the 4200 TapeStation System and RNA quantity was measured using the Qubit 2.0 Fluorometer. RINs ranged from 1.7 – 9.3 with a mean of 7.6. Sequencing libraries were prepared using NEBNext rRNA Depletion Kit v2 and NEBNext Ultra II Directional RNA Library Prep Kit for Illumina following the manufacturer’s directions. Library quality was evaluated using the 4200 TapeStation and library quantity was measured using the Qubit 2.0 Fluorometer. Samples were sequenced using Illumina NovaSeq with a median of 30 million reads.

### Transcriptional profiling

Low quality RNA-seq reads and adapter sequences were removed with timmomatic 0.39(82). The remaining reads were mapped to the T2T-CMH13v2.0 reference genome using STAR 2.5.3a(83) and read counts were calculated using featureCounts from Subread1.6.4(84). Each timepoint was analyzed individually. Genes with low read counts (genes that did not have at least 10 reads in 5 samples) were removed from analysis. Raw read counts were normalized using RUVg(85) with the 5000 most stably expressed genes between all samples at each time point. Differentially expressed genes were identified using DESeq2(86). To be differentially expressed, the gene had to have |log_2_FC| > 1 and adjusted p-value < 0.05.

### Gene set enrichment analysis, CFU confounder analysis, and gene list analysis

Entrez IDs were assigned using AnnotationDBI(87). Reactome(88) pathway enrichment was performed on DEGs with clusterProfiler(89–91) compareCluster and enrichPathway with a Benjamini-Hochberg (BH) adjusted-pvalue cut off of 0.05. MaAsLin2(30) was used to conduct confounder analyses with TSS normalization and Log transformation for each microbe individually. A gene was considered confounded if the adjusted P-value < 0.05.

Gene lists were manually curated from literature(32, 40), MSigDB Hallmark gene sets(33, 41), Hugo Gene Nomenclature Committee (HGNC)(31), and KEGG(42, 43). The gene lists were then filtered for only genes that were responsive to microbial colonization as determined by hierarchical clustering(92) and log_2_ fold change. Gene list scores were calculated as the median log_2_ fold change from the filtered gene lists. Genes that were not present in at least 2/3 time points (due to being excluded for low read counts) were excluded from gene list heatmap and gene scores. Gene score deviation was calculated through 1,000 bootstrapping iterations, and the 95% confidence intervals were calculated using the bias-corrected and accelerated (BCa) bootstrap method.

### Alternative splicing analysis

The RNA-seq library from 48 hours was re-sequenced and combined with the original sequencing run for greater sequencing depth. Initial processing and mapping to the reference genome was performed as previously described. The only difference was the addition of the STAR parameter --twopassMode Basic. Transcript isoforms were predicted using rMATS-turbo(93) (parameters included --variable-read-length, and --allow-clipping) and delta percent spliced in (ΔPSI) was calculated between each microbial treatment and the donor associated vehicle control from junction reads only (JC). Splicing events were considered significant if the adjusted P-value < 0.05, the |ΔPSI| > 10%, and had an average of at least 5 supporting reads for each condition. For DSG and DEGs comparisons, DEGs were determined as previously described using the alternative splicing dataset to negate confounding effects of read depth. Samples had a median of 112 million reads. K. aerogenes Donors 2 and 5 were excluded from analysis due to low read counts.

Functional domain prediction was performed using SpliceDecoder(56), a computational pipeline that predicts the functional impact of alternative splicing (AS) events. Briefly, first SpliceDecoder searches the annotation GTF file used for rMATS to identify transcripts that perfectly match each splicing event’s inclusion or skipping isoform (referred to as template transcripts). These are then used a references to generate simulated transcripts representing the alternative splicing variants. Second, to map functional domains to each transcript, SpliceDecoder uses UniProt GRCh38(94)-based domain annotations. Since the splicing events were analyzed using T2T-CMH13v2.0 genome, we converted the template and simulated transcript coordinates to GRCh38 using hs1ToHg38.over.chain with UCSC liftOver(95), and mapped domain annotations using BEDTools intersect(95). Third, SpliceDecoder compares domain content between the template and simulated transcripts and reports differences in known protein domains, coding sequences, and UTRs. AlphaFold(96) was used to predict protein folding using default parameters.

### Data analysis and visualization

Data was analyzed and visualized using the following R packages: ComplexHeatmap(97), circlize(98), stringi(99), vroom(100), ggpubr(101), ggplot2(102), tidyverse(103), paletteer(104), and rstatix(105).

## Acknowledgments

We also thank the Genome Technologies service and cyberinfrastructure high performance computing resources at The Jackson Laboratory (JAX). These shared services were supported in part by the JAX Cancer Center (P30 CA034196). Figures were created using Adobe Illustrator and BioRender.

This work was primarily supported by the National Institute of Health Award U19AI142733. JO was additionally supported by 1DP2GM126893-01, 1 R01 AR078634-01 and 5 R21 AR075174-02. MH was supported by T32HG010463.

## Supplementary Figures

**Figure S1.**
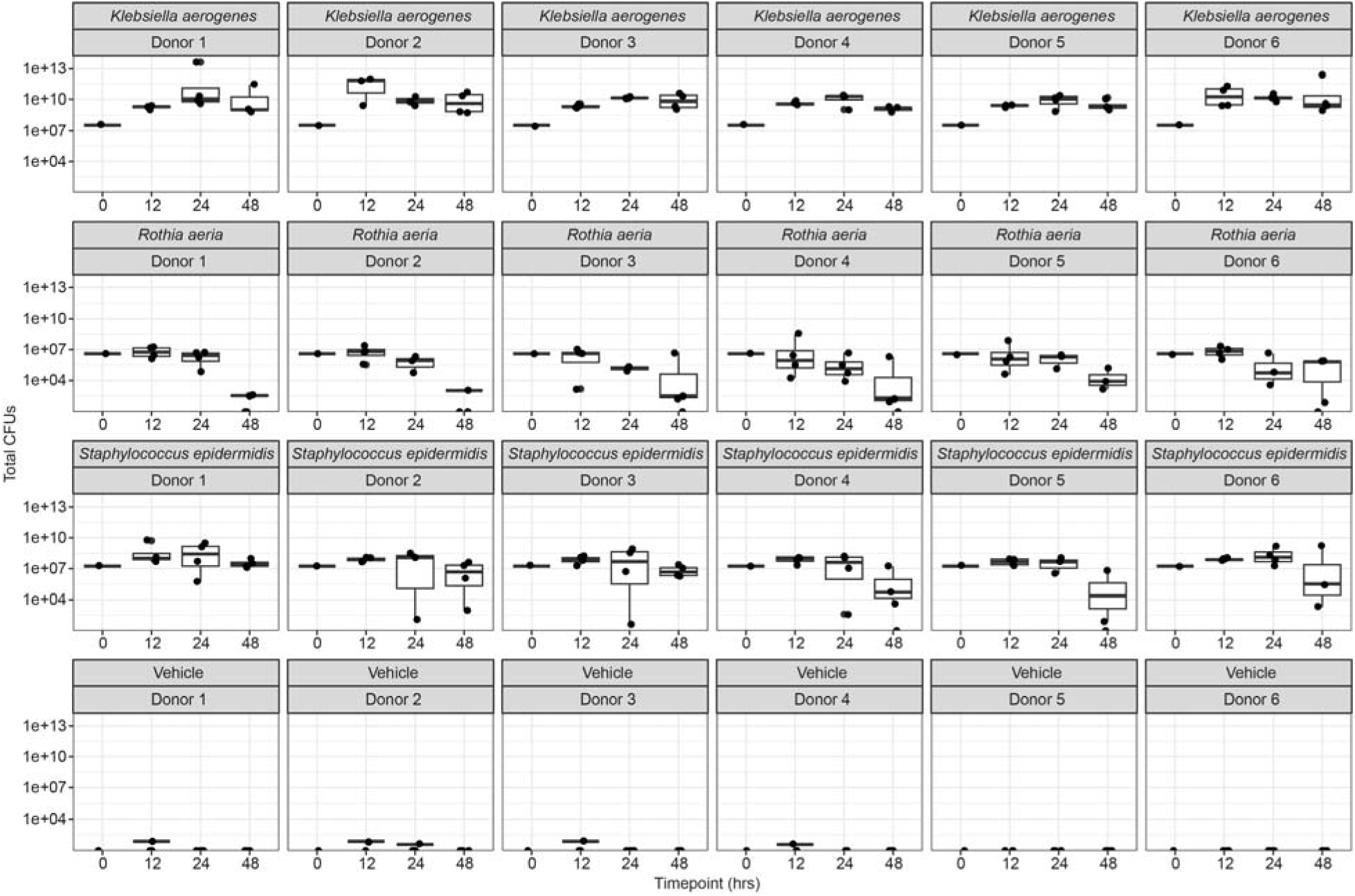
Microbial CFUs of inoculum and at each endpoint. At time of dosing (0 hours) and each time point (12 hours, 24 hours, and 48 hours), the bacterial inoculum or apical wash was serially diluted and plated on tryptic soy agar (TSA) to quantify colony forming units (CFUs). For boxplots, box middle represents the median, box edges represent 25^th^ and 75^th^ quartiles, and outlier values are separate points.

**Figure S2.**
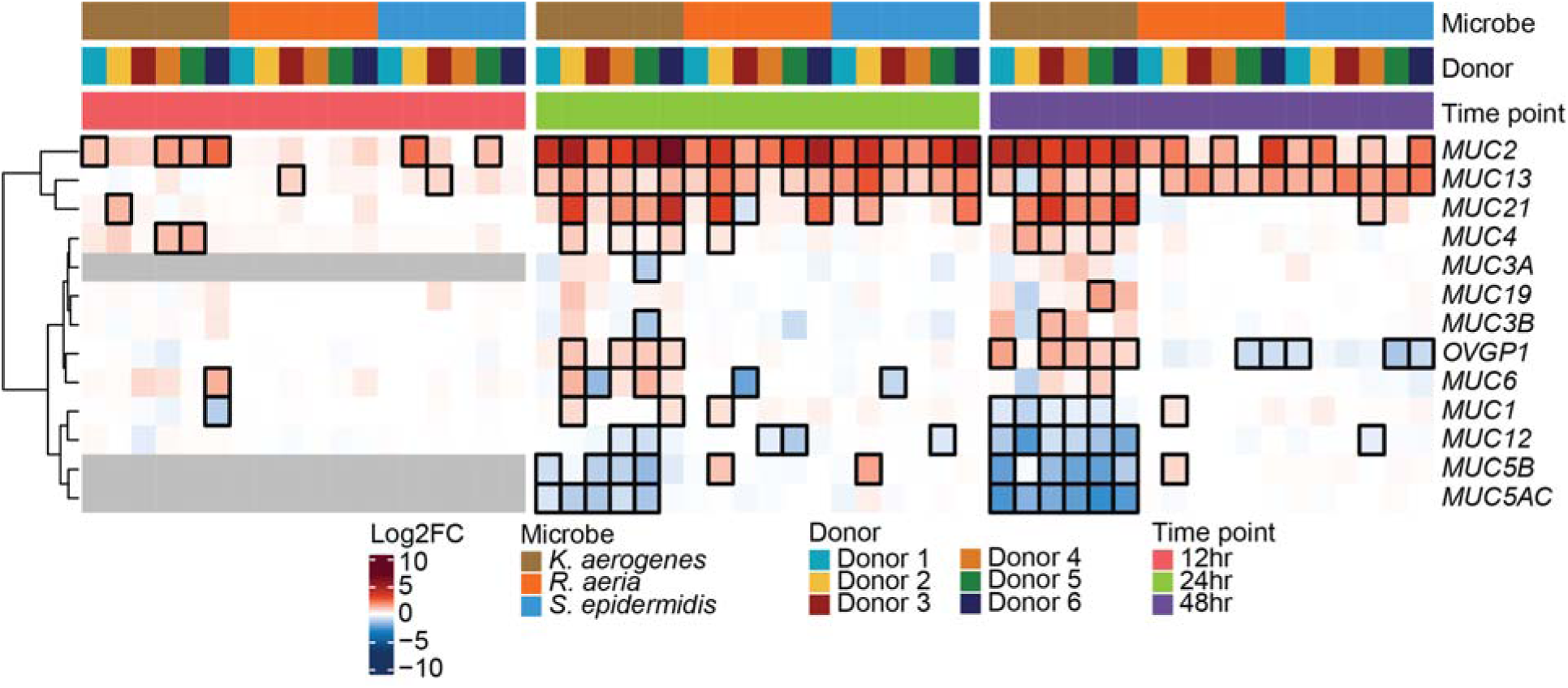
Changes in gene expression of mucin-related genes. Heatmap of mucin-related genes, which comprise mucus. Each row represents a different gene and each column a different sample. Columns were hierarchically clustered. Each column was annotated by color with the sample’s microbial treatment, donor cells, and time point. Cells were colored by the log_2_ fold change (log_2_FC) relative to the appropriate vehicle control. Outlined boxes have an adjusted P-value < 0.05. Gray boxes represent genes that were filtered out due to low gene counts at a specific time point. Genes that were not present in at least 2/3 time points were removed.

**Figure S3.**
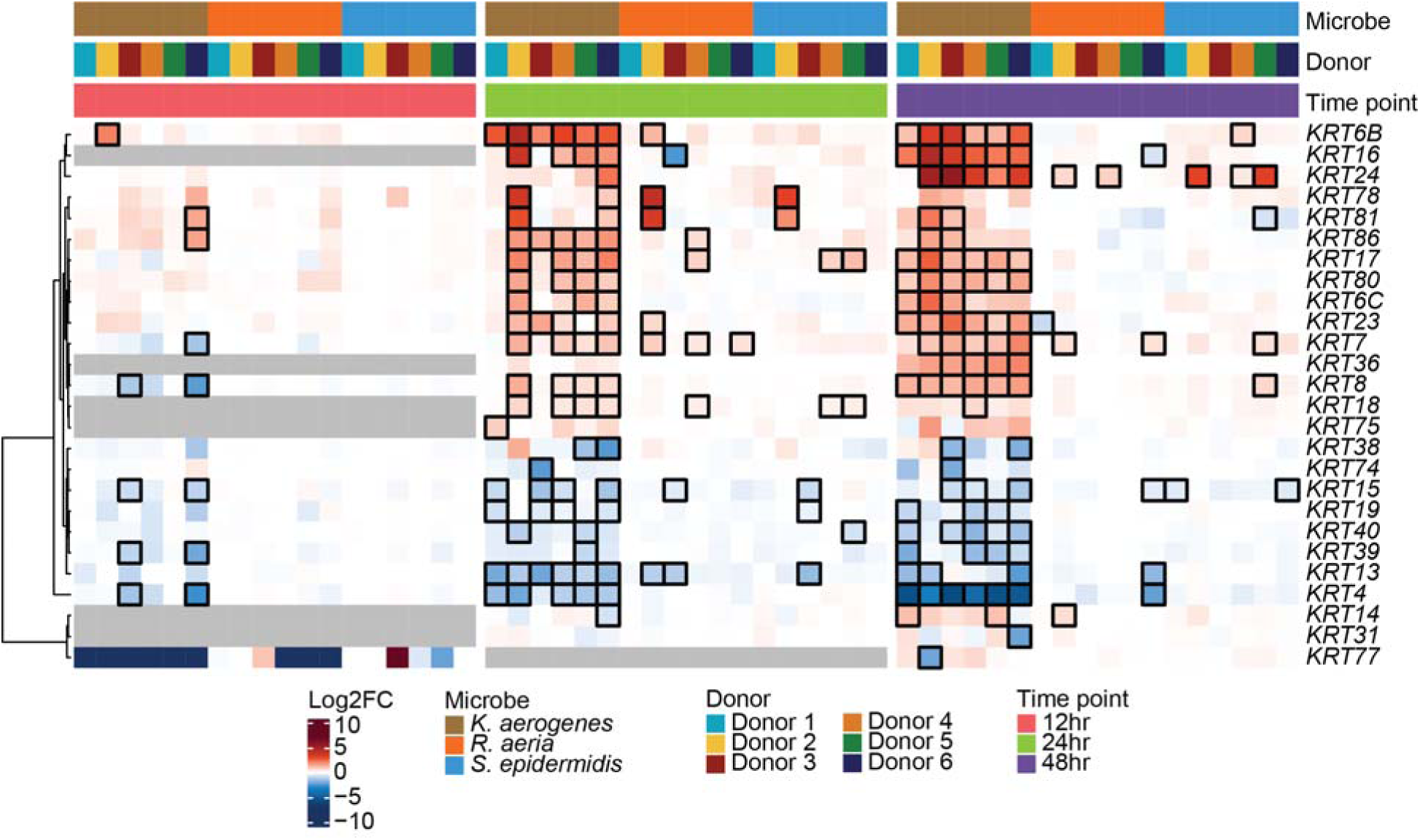
Changes in gene expression of keratin genes. Heatmap of keratin genes, which provide structural integrity. Each row represents a different gene and each column a different sample. Columns were hierarchically clustered. Each column was annotated by color with the sample’s microbial treatment, donor cells, and time point. Cells were colored by the log_2_ fold change (log_2_FC) relative to the appropriate vehicle control. Outlined boxes have an adjusted P-value < 0.05. Gray boxes represent genes that were filtered out due to low gene counts at a specific time point. Genes that were not present in at least 2/3 time points were removed.

**Figure S4.**
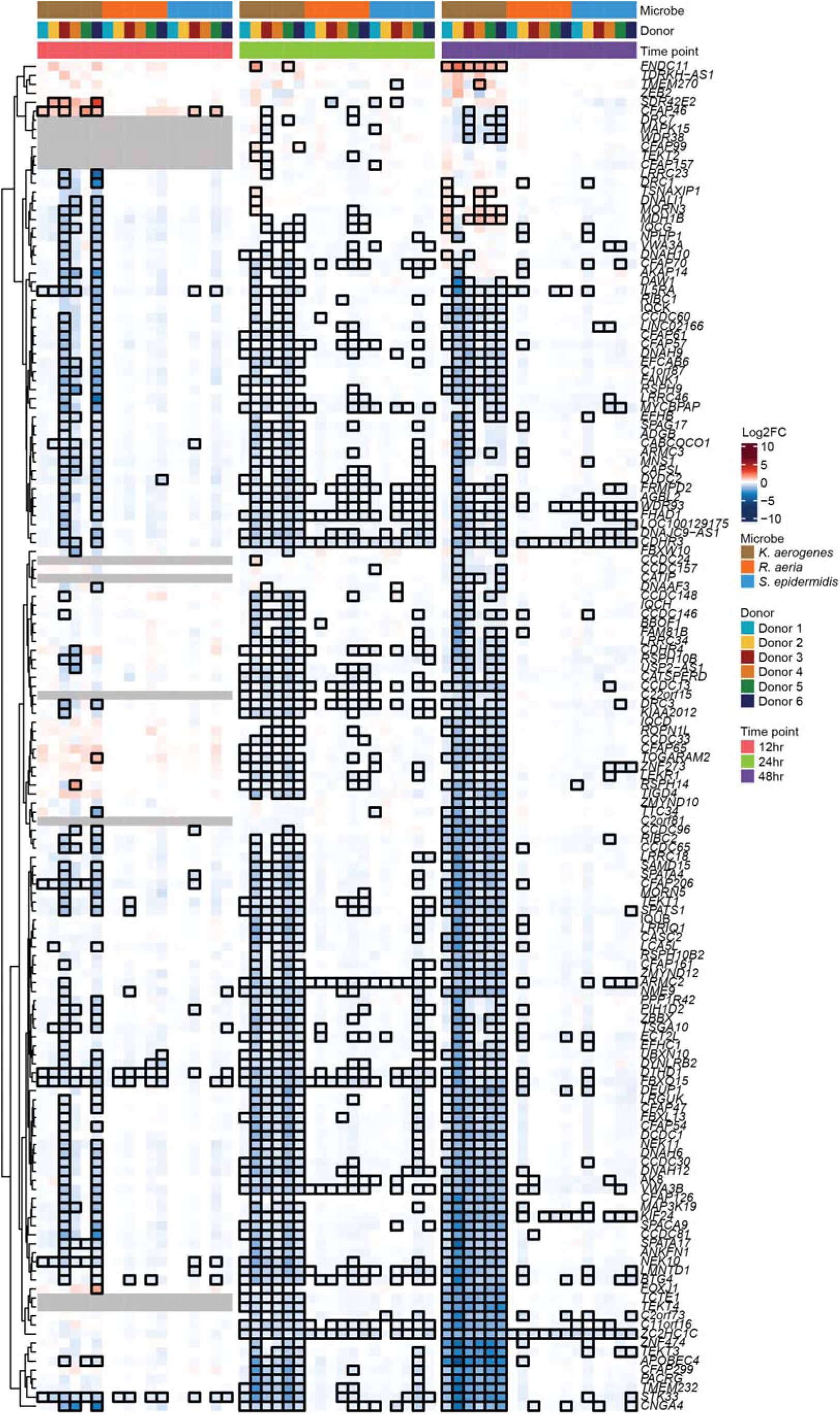
Changes in gene expression of ciliary genes. Heatmap of ciliary genes. Each row represents a different gene and each column a different sample. Columns were hierarchically clustered. Each column was annotated by color with the sample’s microbial treatment, donor cells, and time point. Cells were colored by the log_2_ fold change (log_2_FC) relative to the appropriate vehicle control. Outlined boxes have an adjusted P-value < 0.05. Gray boxes represent genes that were filtered out due to low gene counts at a specific time point. Genes that were not present in at least 2/3 time points were removed.

**Figure S5.**
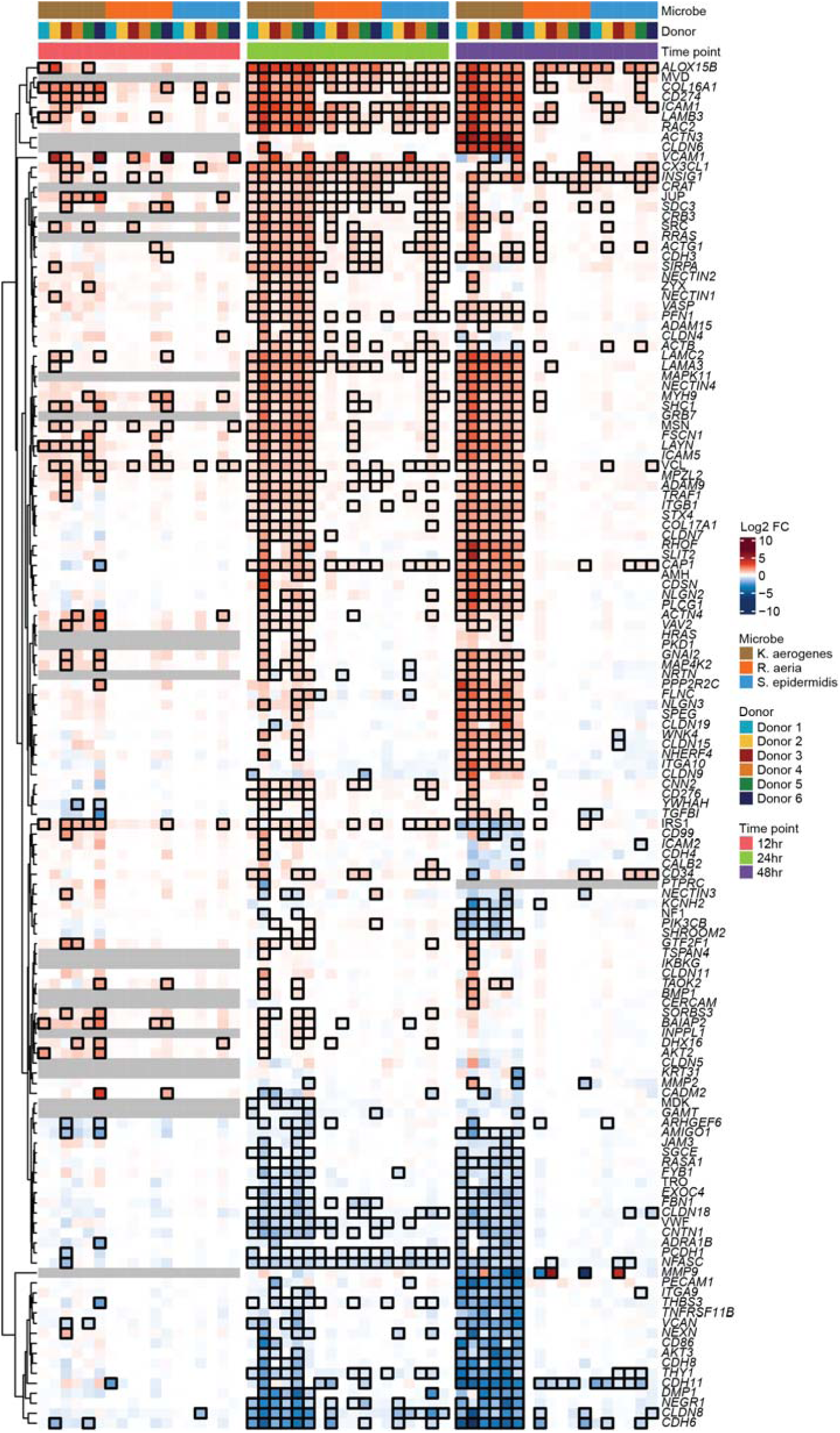
Changes in gene expression of apical junction genes. Heatmap of apical junction genes. Each row represents a different gene and each column a different sample. Columns were hierarchically clustered. Each column was annotated by color with the sample’s microbial treatment, donor cells, and time point. Cells were colored by the log_2_ fold change (log_2_FC) relative to the appropriate vehicle control. Outlined boxes have an adjusted P-value < 0.05. Gray boxes represent genes that were filtered out due to low gene counts at a specific time point. Genes that were not present in at least 2/3 time points were removed.

**Figure S6.**
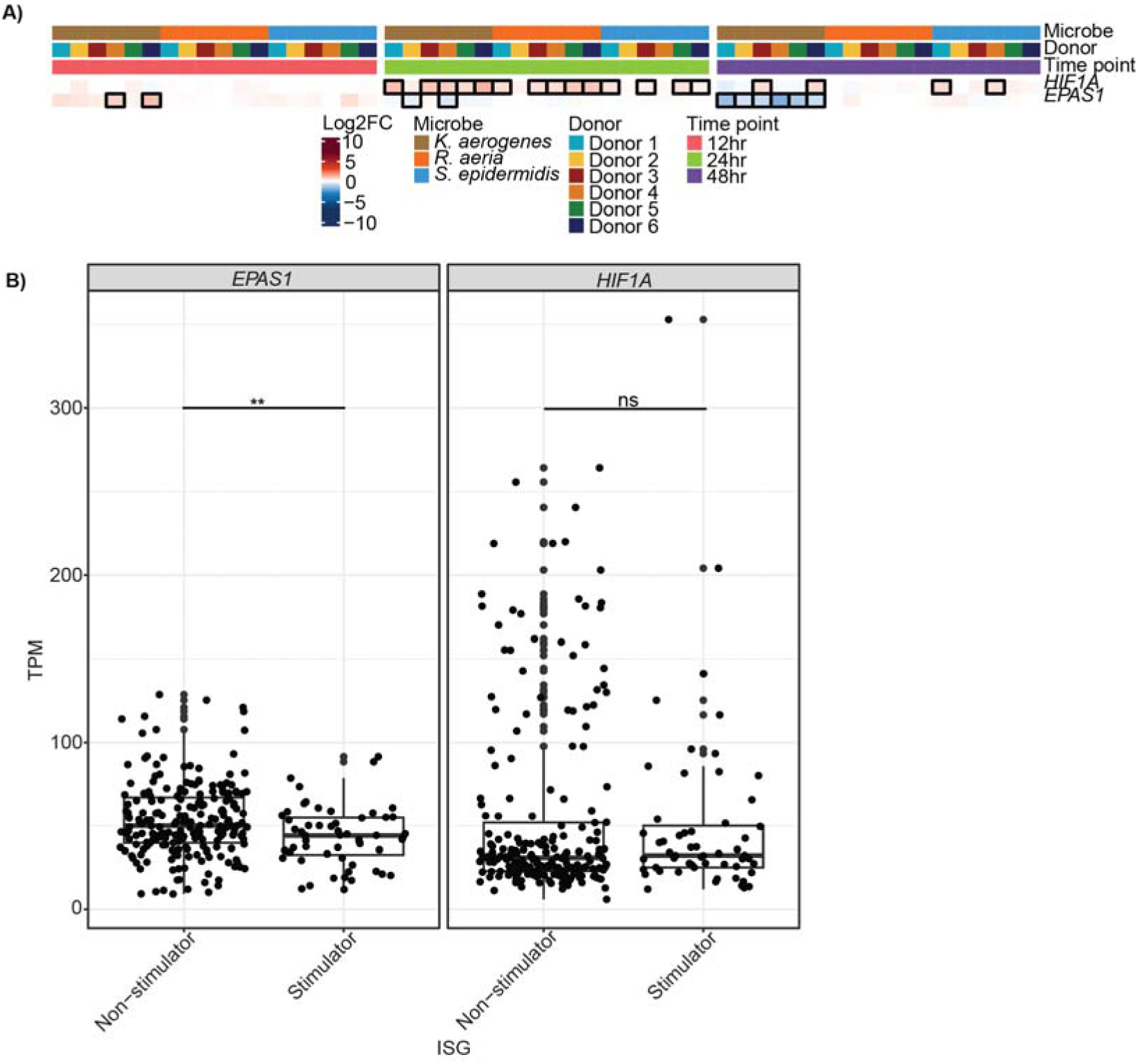
Changes in gene expression of hypoxia genes. **A)** Heatmap of *HIF-1*α and *EPAS1* (HIF-2α), the transcription factors activated by hypoxia. Each row depicts a different gene and each column a different sample. Samples are color annotated according to the donor, microbial treatment, and time point. Rows are hierarchically clustered. Cells are colored based on the log_2_ fold change. Outlined cells represent an FDR-adjusted P-value < 0.05. **B)** Boxplots of TPM (transcripts per million) for *HIF-1*α and *EPAS1*. Comparison using Wilcoxon test. For relevant plots, * represented P-value < 0.05, ** P-value < 0.01, *** P-value < 0.001, and ns = non-significant. For boxplots, box middle represents the median, box edges represent 25^th^ and 75^th^ quartiles, and outlier values are separate points.

**Figure S7.**
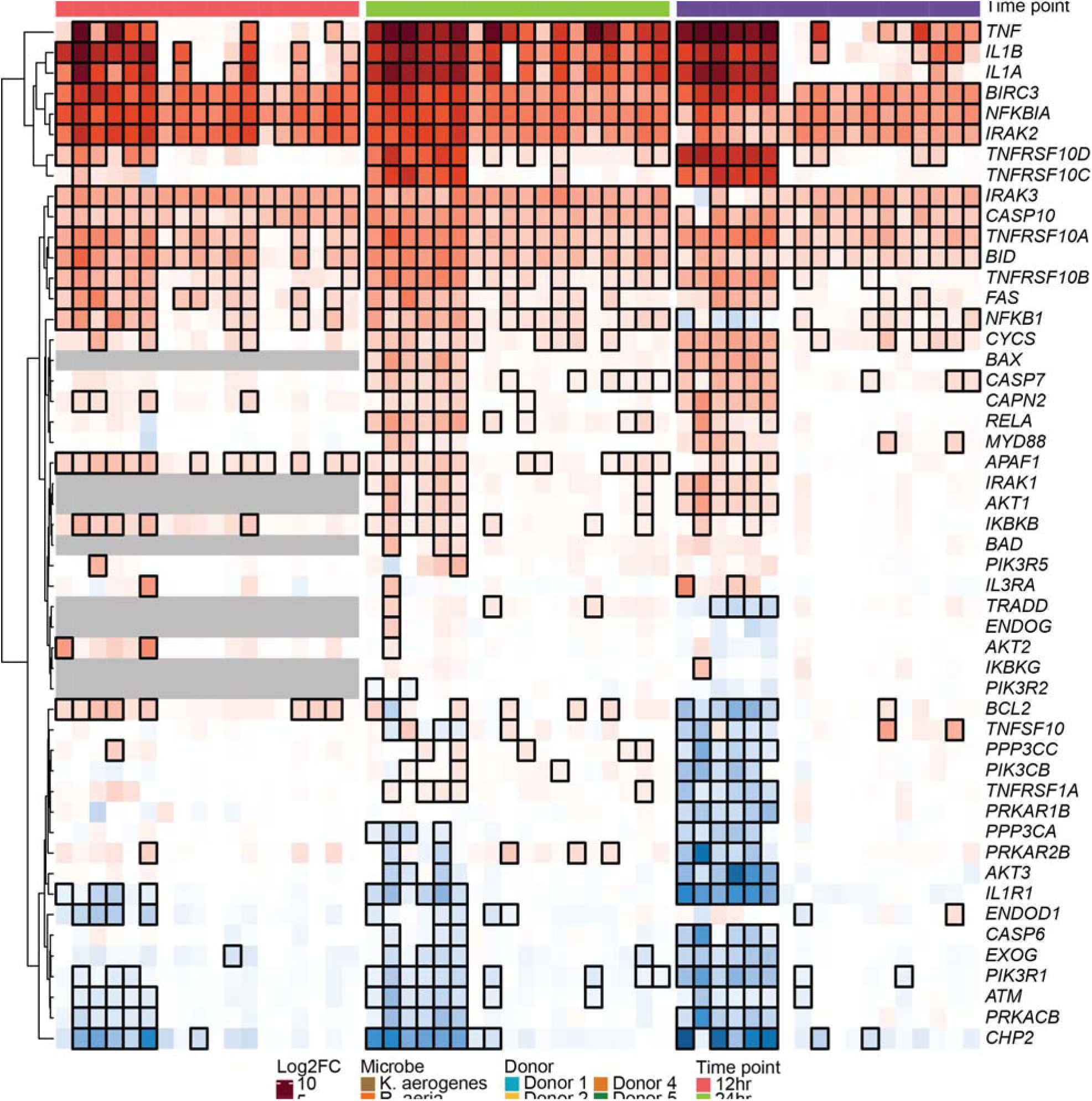
Changes in gene expression of apoptosis-related genes. Heatmap of apoptosis-related genes. Each row represents a different gene and each column a different sample. Columns were hierarchically clustered. Each column was annotated by color with the sample’s microbial treatment, donor cells, and time point. Cells were colored by the log_2_ fold change (log_2_FC) relative to the appropriate vehicle control. Outlined boxes have an adjusted P-value < 0.05. Gray boxes represent genes that were filtered out due to low gene counts at a specific time point. Genes that were not present in at least 2/3 time points were removed.

**Figure S8.**
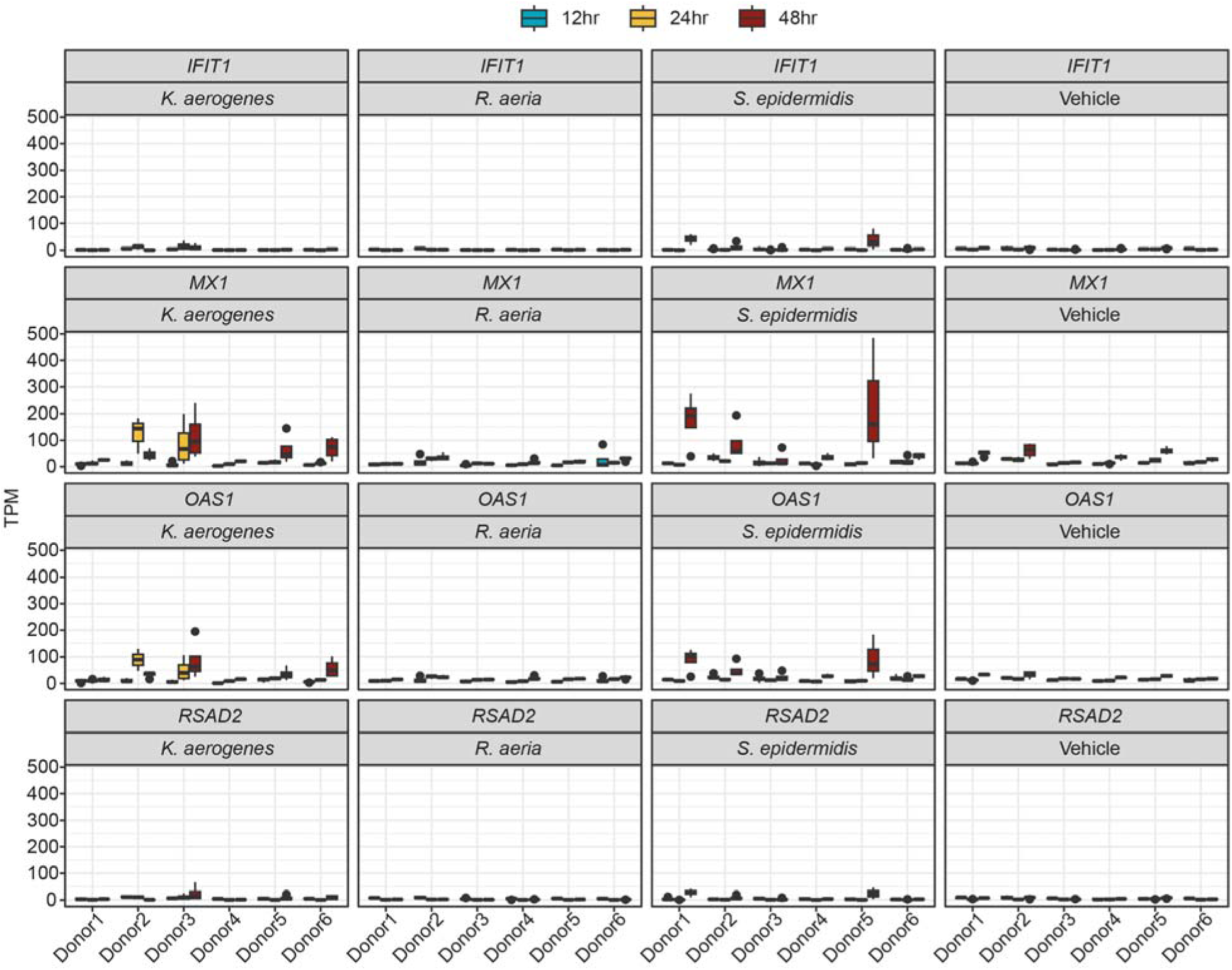
Gene expression of ISGs following microbial treatment versus vehicle. Boxplot of TPM (normalized gene counts) for select ISGs in each microbial treatment and vehicle control. Box middle represents the median, box edges represent 25^th^ and 75^th^ quartiles, and outlier values are separate points. For boxplots, box middle represents the median, box edges represent 25^th^ and 75^th^ quartiles, and outlier values are separate points.

**Figure S9.**
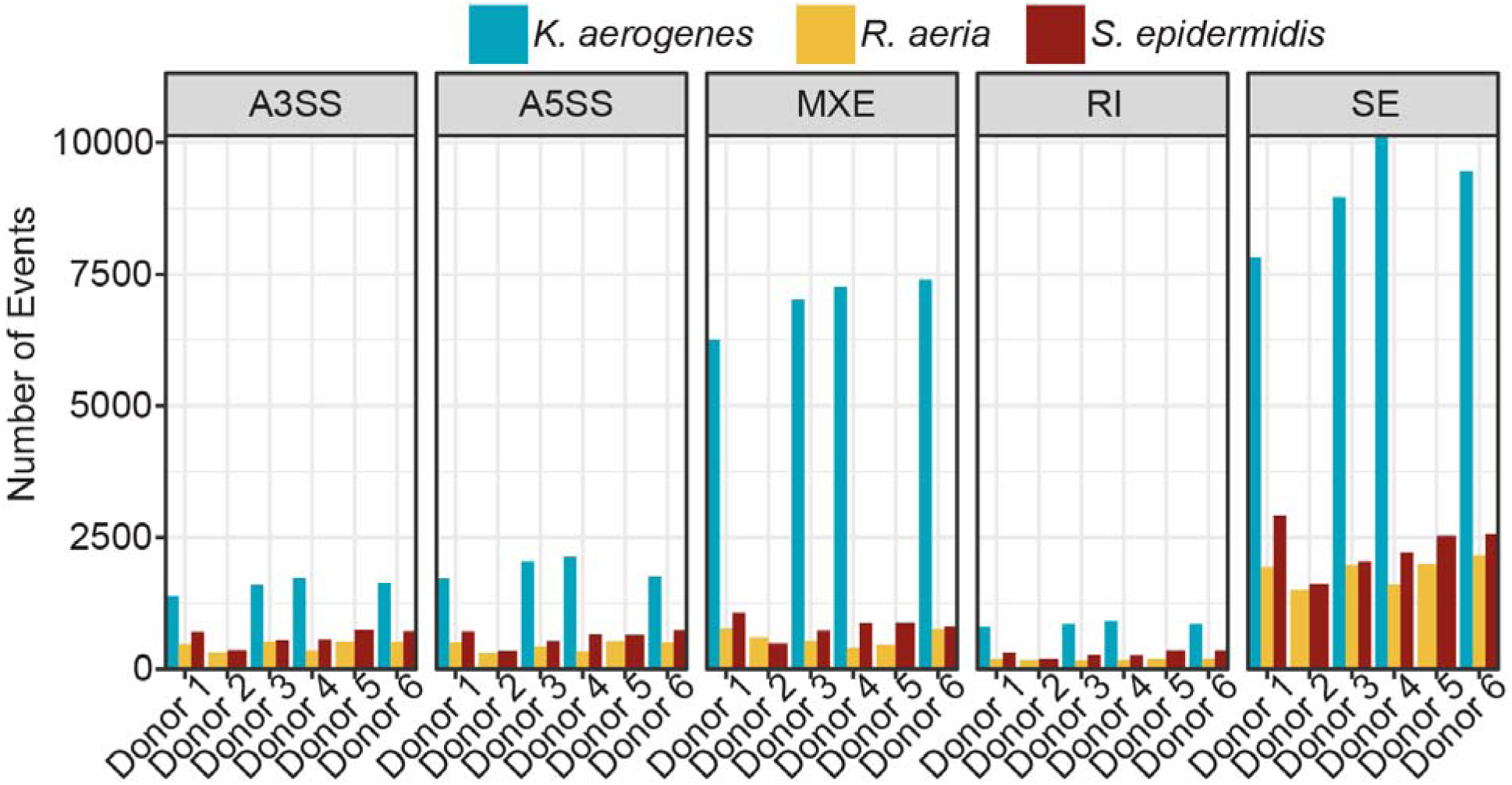
Distribution of differential splicing events. Plot of the number of statistically significant differential splicing events (|ΔPSI| > 0.1 and FDR < 0.05) for each microbe and donor, shown per event type (A3SS: alternative 3’ splice site, A5SS: alternative 5’ splice site, MXE: mutually exclusive exons, RI: retained intron, and SE: skipped exon also called cassette exons).

**Figure S10.**
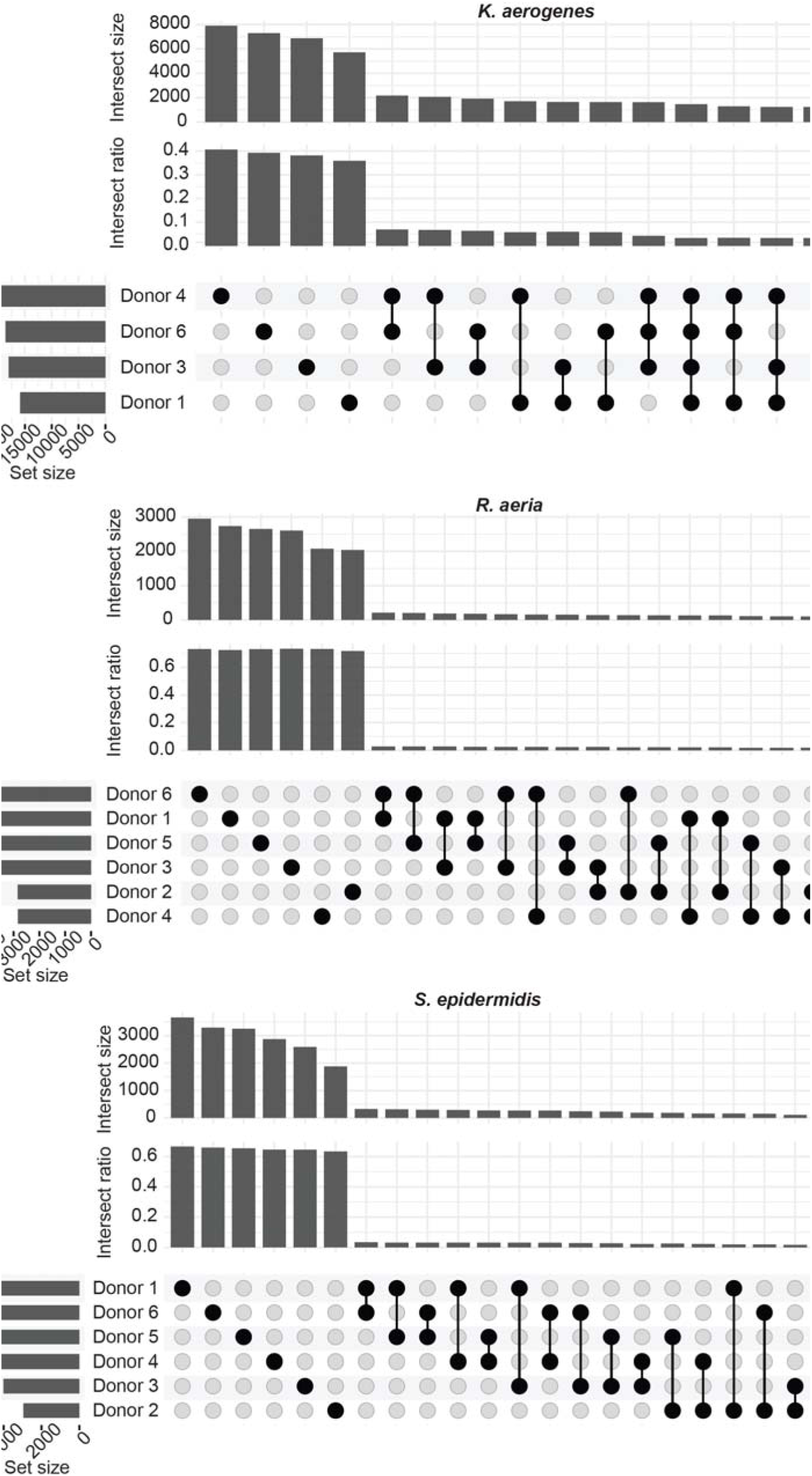
Differentially splicing events shared between donors for each microbe. Upset plot of splicing events depicting how many events are conserved between donors, shown for each microbe: *K. aerogenes* (**A**), *R. aeria* (**B**), and *S. epidermidis* (**C**). Set size represents the total number of splicing events. Donors are sorted by decreasing set size. Intersect size represents the number of splicing events shared between the indicated donors (black points connected by lines). Only intersections with a size of at least 50 are displayed. Intersect ratio normalizes the intersect size by union size (all unique elements across the sets).

**Figure S11.**
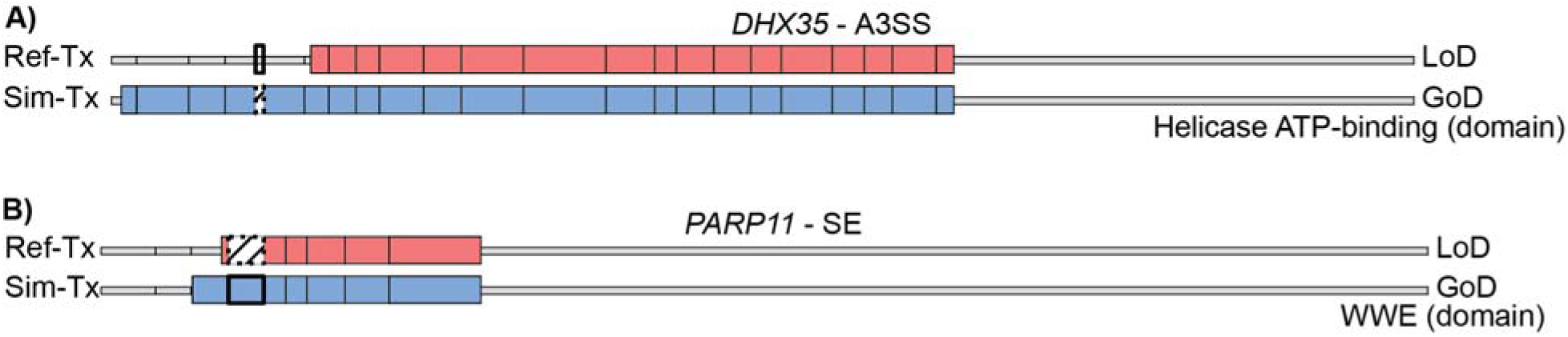
Predicted domain changes to *DHX35* and *PARP11*. Schematic representation of the *DHX35* and *PARP11* transcripts 5’ to 3’, with boxes representing the coding exons and lines representing the UTR regions. Both the reference transcript (Ref-Tx; red) and transcript with the simulated splicing event (Sim-Tx; blue) are shown for each of the two splicing events. The affected domain is labeled. **A)** Usage of an alternative 3’ splice site (A3SS) in *DHX35* resulting in a transcript with a gain of domain (GoD, blue) compared to the reference transcript with the canonical 3’ splice site (red). **B)** A cassette exon in *PARP11* leading to a transcript with a gain of domain (GoD, blue) compared to the transcript without the exon (red).

**Figure S12.**
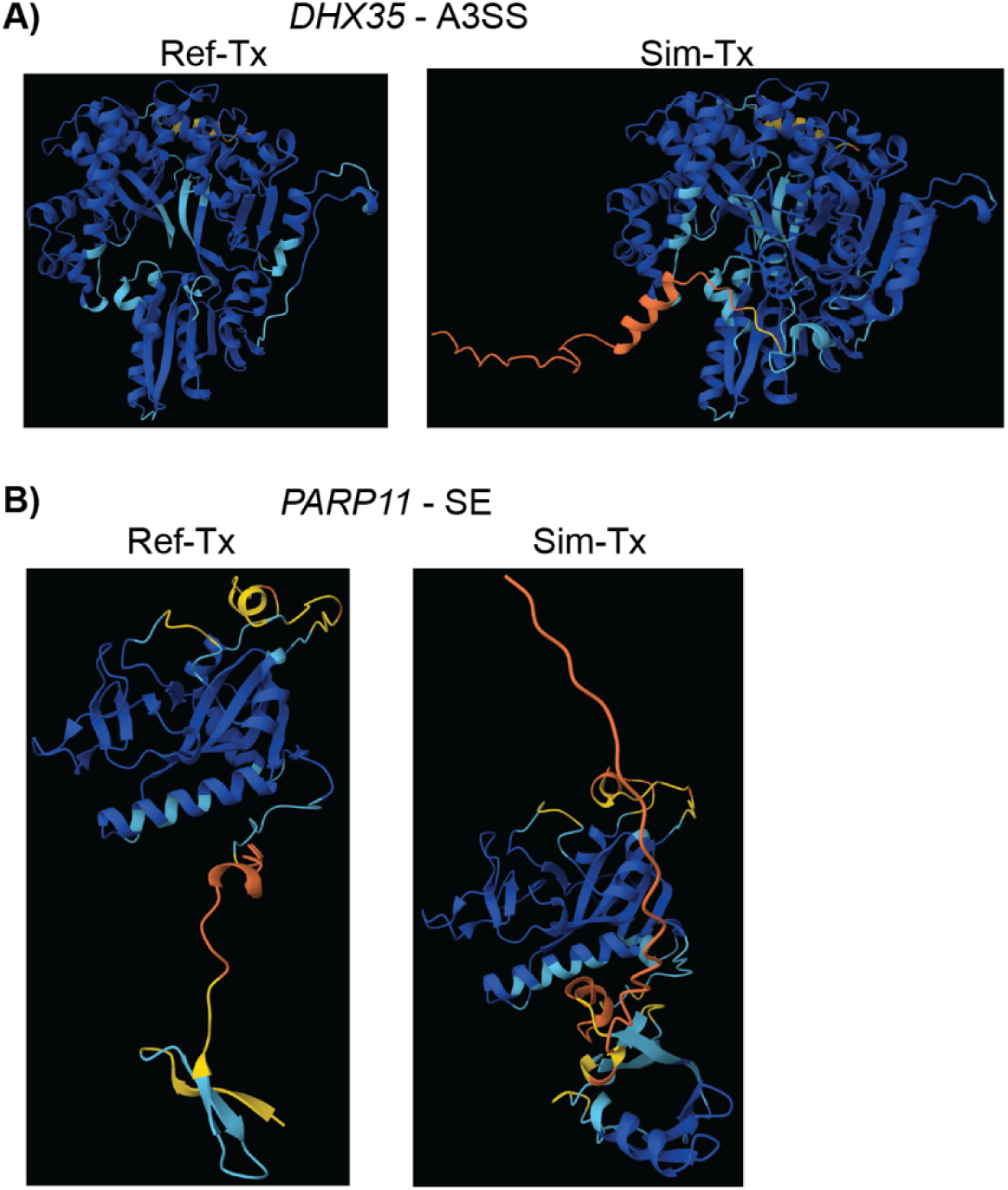
Predicted protein folding of *DHX35* and *PARP11*. Schematic representation of the *DHX35* and *PARP11* proteins where lines represent unstructured regions, arrows represent beta-pleated sheets, and spirals represent alpha-helices. The proteins are colored based on the prediction accuracy (pLDDT), where darker blue represents higher confidence and darker orange lower confidence. Both the reference transcript (Ref-Tx) and transcript with the simulated splicing event (Sim-Tx) are shown for each of the two splicing events. **A)** Usage of an alternative 3’ splice site (A3SS) in *DHX35*. **B)** Inclusion of a cassette exon in *PARP11*.

## Supplementary Tables

**Table S1**: List of donors who provided epithelial cells. For each sample, provided is their age, sex, and race.

**Table S2**: Transepithelial electrical resistance (TEER) measurements collected during ALI maturation for each donor.

**Table S3**: CFUs (colony forming units, a measure of live bacteria) from the inoculum and washed off of the ALI at each harvest for each microbial treatment/vehicle control.

**Table S4**: Number of differentially expressed genes (DEGs) that were upregulated and downregulated in response to each microbial treatment.

**Table S5a**: Reactome pathway analysis of genes differentially expressed between donors in response to microbial treatment following 12 hours of colonization.

**Table S5b**: Reactome pathway analysis of genes differentially expressed between donors in response to microbial treatment following 24 hours of colonization.

**Table S5c**: Reactome pathway analysis of genes differentially expressed between donors in response to microbial treatment following 48 hours of colonization.

**Table S6**: The gene lists used.

**Table S7**: The gene scores calculated for each sample for each gene list.

**Table S8**: Number of differential splicing events (FDR < 0.05 and absolute PSI > 0.1) for each microbial treatment. A3SS = alternative 3’ splice site, A5SS = alternative 5’ splice site, MXE = mutually exclusive exons, RI = retained intron, SE = skipped/cassette exon.

**Table S9**: Number of differentially spliced genes, differentially expressed genes (identified from the deeper read depth), and genes that were differentially spliced and differentially expressed.

**Table S10**: Reactome pathway analysis of differentially spliced genes.

**Table S11**: Number of functional domain changes for each microbial treatment. GoD = gain of domain, LoD = loss of domain, NMD = nonsense-mediated decay, other_region = other functional change, no_change = no functional change.

